# Modeling dynamic oxygen permeability as a mechanism to mitigate oxygen-induced stresses on photosynthesis and N_2_ fixation in marine *Trichodesmium*

**DOI:** 10.1101/2025.02.13.638019

**Authors:** Weicheng Luo, Keisuke Inomura, Ondřej Prášil, Meri Eichner, Ya-Wei Luo

**Affiliations:** State Key Laboratory of Marine Environmental Science and College of Ocean and Earth Sciences, Xiamen University, Xiamen 361102, China; Centre Algatech, Institute of Microbiology of the Czech Academy of Sciences, Třeboň 37901, Czech Republic; Graduate School of Oceanography, University of Rhode Island, Narragansett, RI 02882, USA

**Keywords:** *Trichodesmium*, dynamic oxygen permeability, nitrogen fixation, photorespiration, respiratory protection

## Abstract

*Trichodesmium*, the predominant marine diazotrophic cyanobacterium, concurrently performs nitrogen (N_2_) fixation and photosynthesis, the latter of which produces oxygen (O_2_) that inhibits N_2_ fixation. Hopanoid lipids in *Trichodesmium* may play a role in dynamically regulating membrane permeability to O_2_, potentially alleviating O_2_ stress on N_2_ fixation. However, the physiological impacts of this dynamic permeability are not well understood. We developed a model showing that dynamically modulating membrane O_2_ permeability can enhance N_2_ fixation and growth of *Trichodesmium* by over 50%. High O_2_ permeability (1.5×10^-4^ of O_2_ diffusivity in seawater) during strong photosynthesis accelerates O_2_ exhaust, reducing energy-consuming photorespiration by ∼40%, while low O_2_ permeability (1.0×10^-5^ diffusivity) during active N_2_ fixation minimizes O_2_ stress on N_2_ fixation. Together, these mechanisms increase the carbon and iron use efficiencies by ∼70%. Our study provides a mechanistic and quantitative framework for how dynamic O_2_ permeability benefits *Trichodesmium*, offering insights potentially applicable to other diazotrophs.

**IMPORTANCE:** *Trichodesmium* is a key player in marine N_2_ fixation, essential for oceanic productivity and global biogeochemical cycles. However, a significant challenge arises from the concurrent photosynthetic production of O_2_ during N_2_ fixation, which can inhibit N_2_ fixation and cause energy-wasting photorespiration. We develop a physiological model showing that *Trichodesmium* may dynamically regulate membrane O_2_ permeability to enhance N_2_ fixation and growth. The model suggests two mechanisms: elevated O_2_ permeability during the early daytime of strong photosynthesis accelerates O_2_ exhaust to environment, reducing photorespiration, while reduced O_2_ permeability later limits O_2_ influx from environment, lowering wasteful respiration and maintaining a low intracellular O_2_ level for active N_2_ fixation. These adaptations improve the efficiency of carbon and iron utilization, thereby facilitating N_2_ fixation and growth in *Trichodesmium*. This study sheds light on how *Trichodesmium* and other N_2_-fixing microorganisms can optimize their physiological processes in response to environmental challenges.

**HIGHLIGHTS:**

- We developed a metabolic flux model of *Trichodesmium*, which resolves Dynamic cellular Permeability to O_2_ (DPO_2_).
- DPO_2_ increases N_2_ fixation and growth rates.
- DPO_2_ increases growth efficiency by reducing carbon wasting processes such as photorespiration and respiratory protection.
- DPO_2_, as a result, also increases iron utilization efficiency.

## INTRODUCTION

*Trichodesmium* is a major photoautotrophic contributor to marine nitrogen (N_2_) fixation (e.g., 1-3). *Trichodesmium* faces physiological challenges, such as the decrease in the activity of nitrogenase (the enzyme for N_2_ fixation) upon exposure to oxygen (O_2_) (4-6). Given that *Trichodesmium* simultaneously conducts N_2_ fixation and O_2_-producing photosynthesis during the daytime (e.g., 4, 7), *Trichodesmium* has developed several physiological strategies to cope with this O_2_ stress on nitrogenase and protect N_2_ fixation (8). One of these is the respiratory protection: *Trichodesmium* creates low-O_2_ intracellular environment to realize sufficient N_2_ fixation by temporally segregating photosynthesis and N_2_ fixation and wastefully respiring organic carbon with intracellular O_2_ (9-11). A potentially complementary strategy is the diffusion adjustment; a proper low cell membrane permeability to O_2_ can contribute to forming and maintaining the low-O_2_ window (10, 12, 13).

Notably, during the early light period with high rates of photosynthesis and O_2_ production, low cell permeability to O_2_ could lead to high intracellular O_2_ concentrations, resulting in oxidative stress on photosynthesis as well as “photorespiration”, an energy inefficient consumption of O_2_ (10, 13, 14). Photorespiration is a light-dependent process, which consumes ATP (adenosine triphosphate), NADPH (nicotinamide adenine dinucleotide phosphate hydrogen) and O_2_ and produces CO_2_ (15). This oxygenation reaction is catalyzed by RuBisCO (ribulose-1,5-bisphosphate carboxylase/oxygenase) with RuBP (ribulose-1,5-bisphosphate) and O_2_ as substrates (15). Consequently, the high intracellular O_2_ concentration during early daytime can compete with CO_2_ and inhibits the carboxylation activity of RuBisCO, thus increasing photorespiration and reducing photosynthetic carbon fixation (13, 14, 16). While cyanobacteria are known to operate carbon concentrating mechanisms to increase intracellular CO_2_ concentration and thus shift the CO_2_:O_2_ ratio in favor of carboxylation rather than oxygenation, recent studies have suggested that photorespiration may still play an important role in these organisms (17). Technical difficulties have so-far hindered direct quantification of photorespiration rates in cyanobacteria and given these uncertainties, photorespiration has not been explicitly resolved in previous models of *Trichodesmium* (10, 12, 13).

Due to the importance of O_2_ concentration on the likelihood of photorespiration, the permeability of the cell membrane to O_2_ may impact the occurrence of photorespiration. For example, a high cell permeability to O_2_ could facilitate the rapid diffusion of intracellular O_2_ to extracellular environment, likely decreasing the photorespiration rate during early light period when photosynthesis is strong in *Trichodesmium* (14). This high O_2_ permeability could also enhance the diffusion rate of extracellular O_2_ into the cytoplasm during the low-O_2_ window, thereby elevating the respiratory protection required to consume organic carbon and intracellular O_2_ as an indirect cost for N_2_ fixation (10). On the contrary, a low cell permeability to O_2_ would elevate the photorespiration, but benefit N_2_ fixation. The above scenarios are based on the assumption of diurnally constant cell permeability to O_2_, as employed in previous model studies (10, 12, 13). However, *Trichodesmium* can synthesize hopanoids, intercalated into lipid bilayers of membranes and possibly forming rafts to redistribute hopanoid molecules (18, 19), which might allow diurnal changes in cell permeability to O_2_. The potential physiological implications of this Dynamic cell Permeability to O_2_ (DPO_2_) to *Trichodesmium* remain poorly understood and warrant further investigation.

In this study, we hypothesize that DPO_2_ help to regulate intracellular O_2_ levels and mitigate O_2_-induced stresses in *Trichodesmium*, promoting the efficiency of key enzymes such as RuBisCO and nitrogenase in the context of temporally segregating the activities of the two enzymes. To test the hypothesis, we improved previous models by representing more processes including photorespiration and DPO_2_. The analyses of the model results, along with the comparison to additional model experiments of fixed O_2_ permeability, provided a mechanistic and quantitative understanding of the potential role of DPO_2_ in impacting photorespiration and N_2_ fixation in *Trichodesmium*.

## METHODS

The model in this study was developed by incorporating new representations of photorespiration and DPO_2_ into previous models (10, 20). In the subsequent sections, we present a concise overview of the model schemes. We provide more detailed descriptions, parameter values, intermediate variables and state variables in Supplementary Methods and Tables S1 to S3.

### General model framework

The model (Fig. 1A) simulates key physiological processes in *Trichodesmium* trichome, including photosynthetic electron transfer (PET), carbon fixation, photorespiration and N_2_ fixation over a 12-hour diurnal cycle. These processes are modulated by the dynamic allocation of Fe, ATP and NADPH to different metabolic pathways, as well as by intracellular O_2_ management (Fig. 1A). Two pathways of PET are represented, including linear PET (LPET) that produces ATP and NADPH, and alternative electron transfer (AET) that only generates ATP. ATP and NADPH are used in various processes as described below. N_2_ fixation occurs only when intracellular O_2_ is low. Photorespiration increases with increasing intracellular O_2_ levels.

**FIG 1.**
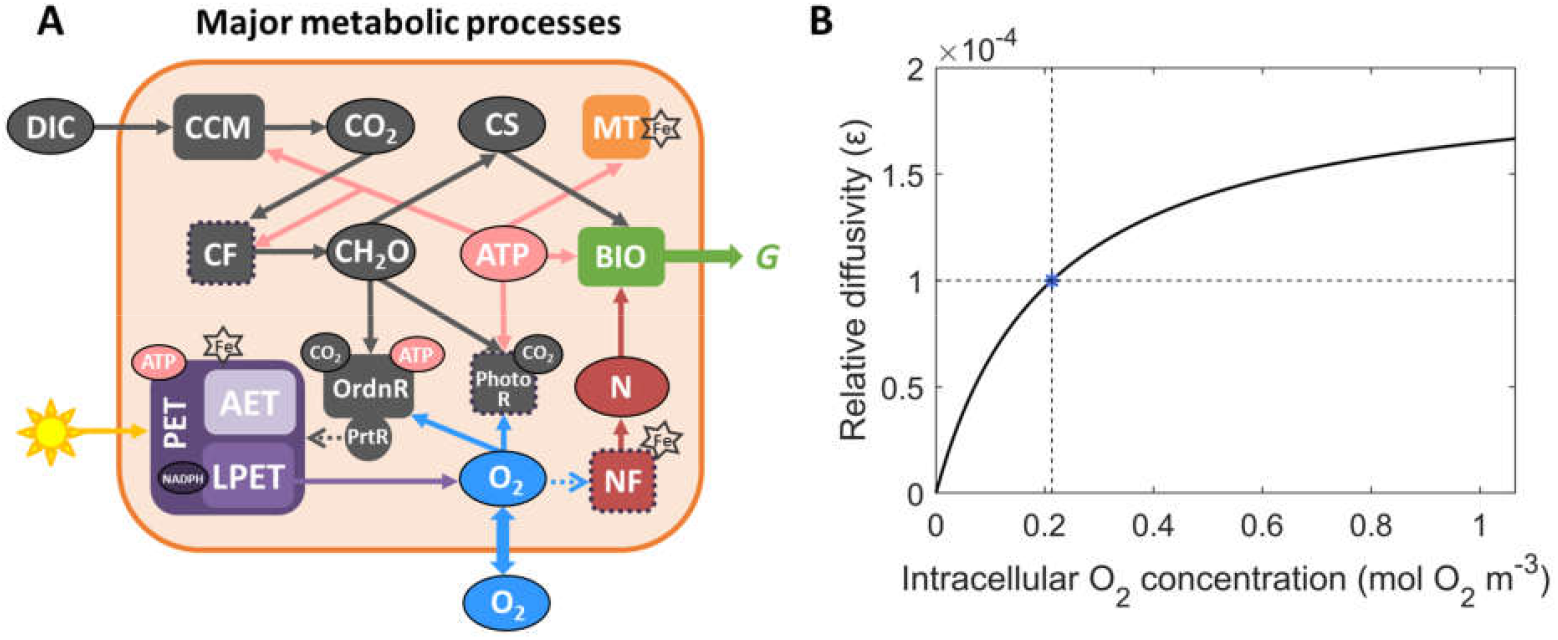
Schematic of the physiological model with potential dynamic O_2_ permeability in *Trichodesmium*. (**A**) Photosynthesis, photorespiration (PhotoR), N_2_ fixation and other key processes simulated within *Trichodesmium* trichome. The pentagrams marked Fe-requiring processes, with fundamental intracellular Fe pools showed in Fig. S1A. Dashed arrows represent inhibition effects. CF: carbon fixation; NF: N_2_ fixation; PrtR: respiratory protection; OrdnR: ordinary respiration; CH_2_O: carbohydrate; CS: carbon skeleton; N: fixed nitrogen; MT: maintenance; BIO: biosynthesis; *G*: growth rate. (**B**) The model parameterizes the dynamic O_2_ permeability by assuming that the intracellular O_2_ concentration regulates the O_2_ diffusion coefficient of the cell membrane relative to that of seawater. Blue asterisk denotes the relative diffusivity of the cell membrane at the half-saturating coefficient of intracellular O_2_ concentration (0.213 mol O_2_ m^-3^).

### Photosynthetic pathways

The photosynthetic pathways followed previous model schemes (10, 20). The total rate of PET is positively regulated by light intensity and the Fe allocated to photosystems. Conversely, it is mitigated by respiratory protection mechanisms (4). The proportion of electrons directed towards linear PET (LPET) and alternative electron transport (AET) is computed at each time step. This is assumed to meet the immediate intracellular requirements for ATP and/or NADPH (10, 20).

### O_2_ production and dynamic permeability

O_2_ is exclusively produced by LPET (21, 22). AET reduces photosynthetic O_2_ production while also supporting ATP production (21-23). Additionally, *Trichodesmium* can perform respiratory protection to wastefully consume organic carbon and intracellular O_2_ and protect N_2_ fixation (4, 10, 12, 24, 25). O_2_ can also physically diffuse between cytoplasm and extracellular environment (Fig. 1A). The direction and rate of O_2_ diffusion depend on the difference between intracellular and extracellular O_2_ concentration, as well as the O_2_ permeability of the cell membrane (26).

In “dynamic-permeability” model case, the membrane O_2_ permeability is assumed to vary diurnally (18). The membrane O_2_ permeability (*ε*), represented as the relative diffusivity to seawater, is paramerized to increase with intracellular O_2_ concentration using a Michaelis-Menten equation (Fig. 1B):

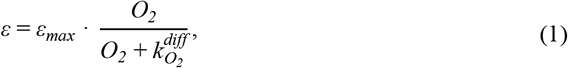

where *ε*_*max*_ is the maximal relative diffusion coefficient of 2.0×10^-4^, and 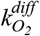 (0.213 mol O_2_ m^-3^), a saturating concentration in seawater at 34 PSU salinity and 25 °C (27), is the half-saturation constant of O_2_ for *ε*.

In addition, to quantitatively assess the physiological roles of DPO_2_, we performed another “fixed-permeability” model case with a fixed *ε* of 1.0×10^-4^.

Given *ε*, the rate of O_2_ diffusion 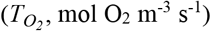 between the intracellular cytoplasm and the extracellular environment is calculated using the scheme proposed by Staal et al. (26) for cylinder-shaped cells:

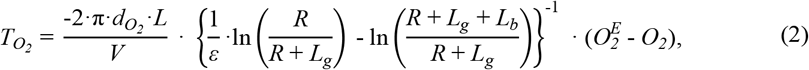

Where 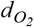 is the O_2_ diffusion coefficient in seawater at 34 PSU and 25 °C, *L* (m) and *V* (m^3^) represent the length and the volume of the trichome, *R* (m) represents the radius of the cytoplasm, *L*_*g*_ (m) denotes the thickness of the cell membrane, *L*_*b*_ (m) denotes the thickness of the boundary layer, 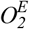 refers to the extracellular far-field O_2_ concentration.

### Photorespiration

Our model additionally represents photorespiration. Photorespiration requires energy using organic carbon and O_2_ as substrates (14, 28). The energy usage by photorespiration consequently reduces the energy availability for carbon and N_2_ fixations. The maximal photorespiration rate [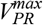, mol C (mol C)^−1^ s^−1^] is computed based on the assumption that produced ATP by PET is fully consumed by photorespiration:

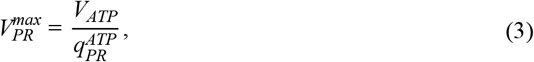

Where 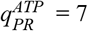 mol ATP (mol C)^-1^ is ATP to C ratio in photorespiration (15).

The rate of photorespiration [*V*_*PR*_, mol C (mol C)^-1^ s^-1^] is also regulated by substrates including intracellular carbohydrate [*CH*_*2*_*O*, mol C (mol C)^-1^] and O_2_ [*O*_*2*_, mol O_2_ m^-3^]:

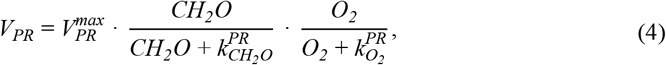

Where 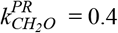 [mol C (mol C)^-1^] and 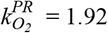 (mol O_2_ m^-3^) are half-saturating coefficients of *CH*_*2*_*O* and *O*_*2*_ for photorespiration (Fig. S1B).

The NADPH, ATP and O_2_ consumption rates of photorespiration [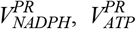, and 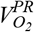 mol NADPH (mol C)^-1^ s^-1^, mol ATP (mol C)^-1^ s^-1^ and mol O_2_ (mol C)^-1^ s^-1^] are:

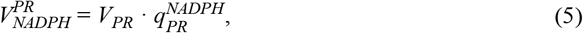

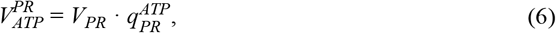

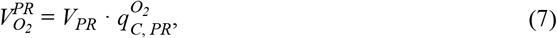

Where 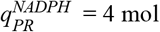 NADPH (mol C)^-1^ and 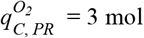 O_2_ (mol C)^-1^ are NADPH to C and O_2_ to C ratios in photorespiration (15).

### N_2_ fixation

N_2_ fixation was calculated according to previous model schemes (10, 20). N_2_ fixation necessitates the utilization of both ATP and NADPH (29, 30). The maximal potential of N_2_ fixation rate [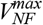, mol N (mol C)^-1^ s^-1^] occurs when the produced ATP and NADPH from PET are completely consumed by N_2_ fixation (10). The rate of N_2_ fixation [*V*_*NF*_, mol N (mol C)^-1^ s^-1^] (*see* Supplementary Methods) is limited by the Fe allocated to nitrogenase [*Fe*_*NF*_, μmol Fe (mol C)^-1^] (31-33) and can be impended by intracellular O_2_, with the rate decreasing upon exposure to O_2_ (34).

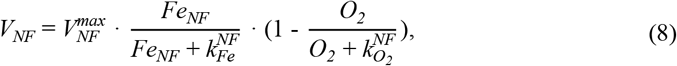

where 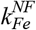 [μmol Fe (mol C)^-1^] and 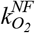 (mol O_2_ m^-3^) are half-saturating coefficients of *Fe*_*NF*_ and *O*_*2*_ for N_2_ fixation.

Respiratory protection also followed previous model schemes (10, 20). Respiratory protection is a mechanism that involves the wasteful respiration of carbohydrates to reduce intracellular O_2_ concentration, thereby supporting N_2_ fixation (12). The rate of respiratory protection increases in response to the demand for N_2_ fixation, while it decreases as intracellular O_2_ level rises (10, 20).

### Carbon fixation

Carbon fixation was computed upon previous models (10, 20) with a minor change by considering photorespiration. Similarly to N_2_ fixation, carbon fixation also relies on the availability of both NADPH and ATP (35). To calculate the carbon fixation rate, the total production of NADPH and ATP at each time step is assumed to be promptly and completely utilized by intracellular processes (10, 20), including photorespiration, CCM, carbon fixation, N_2_ fixation and maintenance (10, 20).

Carbohydrates, which are generated through carbon fixation, stimulate the production of carbon skeletons. However, this production is subsequently downregulated due to the accumulation of these carbon skeletons (*see* Supplementary Methods).

### Intracellular Fe pools and translocation

This part was conducted based on previous model schemes (10, 20). The total intracellular Fe, which encompasses both metabolism and storage (Fig. S1A), is calculated using a previously established scheme (31). Metabolic Fe includes Fe in photosystems, nitrogenase, maintenance and buffer (Fig. S1A). Fe utilized by the photosystems and nitrogenase is from the buffer pool (Fig. S1A). The parameterization of synthesis and decomposition rate of photosystems, as well as the synthesis rate of active nitrogenase and its inactivation, is based on a recent model study featuring diurnally dynamic Fe allocation (20).

Considering that DPO_2_ may contibute to improving the efficiency of Fe in photosystems and nitrogenase, thus regulating intracelluar Fe allocation, model cases were run under various Fe levels for comparison.

### Model parameter values

In both fixed and dynamic O_2_ permeability model cases, four parameters were optimized to maximize the growth rate of *Trichodesmium* (36): including the maximal respiratory protection rate 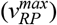, the maximal synthesis 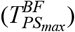 and decomposition 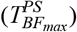 rates of photosystems and the maximal synthesis rate 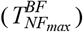 of nitrogenase (Table S1). The optimization was performed employing the global optimizer MultiStart in MATLAB.

Other parameters (Table S2) were either adopted from previous studies or tuned to fit the observed growth rates, N_2_ fixation rates and diurnal Fe in photosystems and nitrogenase from a laboratory culture experiment (32). Given that the experiment was conducted under constant light intensity (90 μmol m^-2^ s^-1^), we adopted the same light intensity with DPO_2_. After tuning model parameters (Table S2), the modeled growth rates (0.28 and 0.45 d^-1^ under low and high Fe, respectively) were well aligned with the observations. Moreover, the model reproduced diurnal patterns of photosystem and nitrogenase (under low and high Fe, for photosystem Fe, *R*^*2*^ = 0.13 and 0.90, respectively; for nitrogenase Fe, *R*^*2*^ = 0.60 and 0.80, respectively) (Fig. S2), supporting the robustness of our model.

Note that the constant light intensity was only used when tuning model parameters. All the model results presented in following were simulated with dynamic light intensity using a sine function over a 12-hour light period (37).

## RESULTS

### Simulated growth rate, carbon and N_2_ fixation rates and O_2_ concentration

The simulations encompassed ten Fe levels (20 pM to 1800 pM) as a previous model study (20). Our model showed a positive correlation between Fe concentration and both N_2_ fixation and growth rates (Fig. S3), consistent with trends from previous culturing experiments of *Trichodesmium* (33). Additionally, model results exhibited that the influence of DPO_2_ on promoting growth rates decreased from 78% to 33% as Fe concentration increased (Fig. S3), indicating that physiological benefits of DPO_2_ in *Trichodesmium* were more pronounced under low Fe conditions. In the following, we focused on analyzing the model results at two levels of dissolved inorganic Fe (40 pM and 1250 pM) that were set in the laboratory experiments (32). Under these two Fe levels, DPO_2_ promoted modeled growth rates of *Trichodesmium* by 61% and 30%, respectively.

Our results revealed that while the dynamic-permeability case exhibited higher growth rates than the fixed-permeability case under both Fe conditions, the gross carbon fixation rates in the dynamic-permeability case were even lower (Fig. 2). This implies that dynamic O_2_ permeability led to an improvement in carbon use efficiency of *Trichodesmium*, defined here as the ratio of net to gross carbon production (Fig. 2). The decrease in the requirement for carbon fixation and the increase in carbon use efficiency is attributed to the lowered photorespiration and the downregulated requirement of respiratory protection (*see* Discussion) (Fig. 2). Furthermore, our simulation results aligned with previous studies (20, 38, 39), demonstrating higher carbon use efficiency under higher Fe conditions.

**FIG 2.**
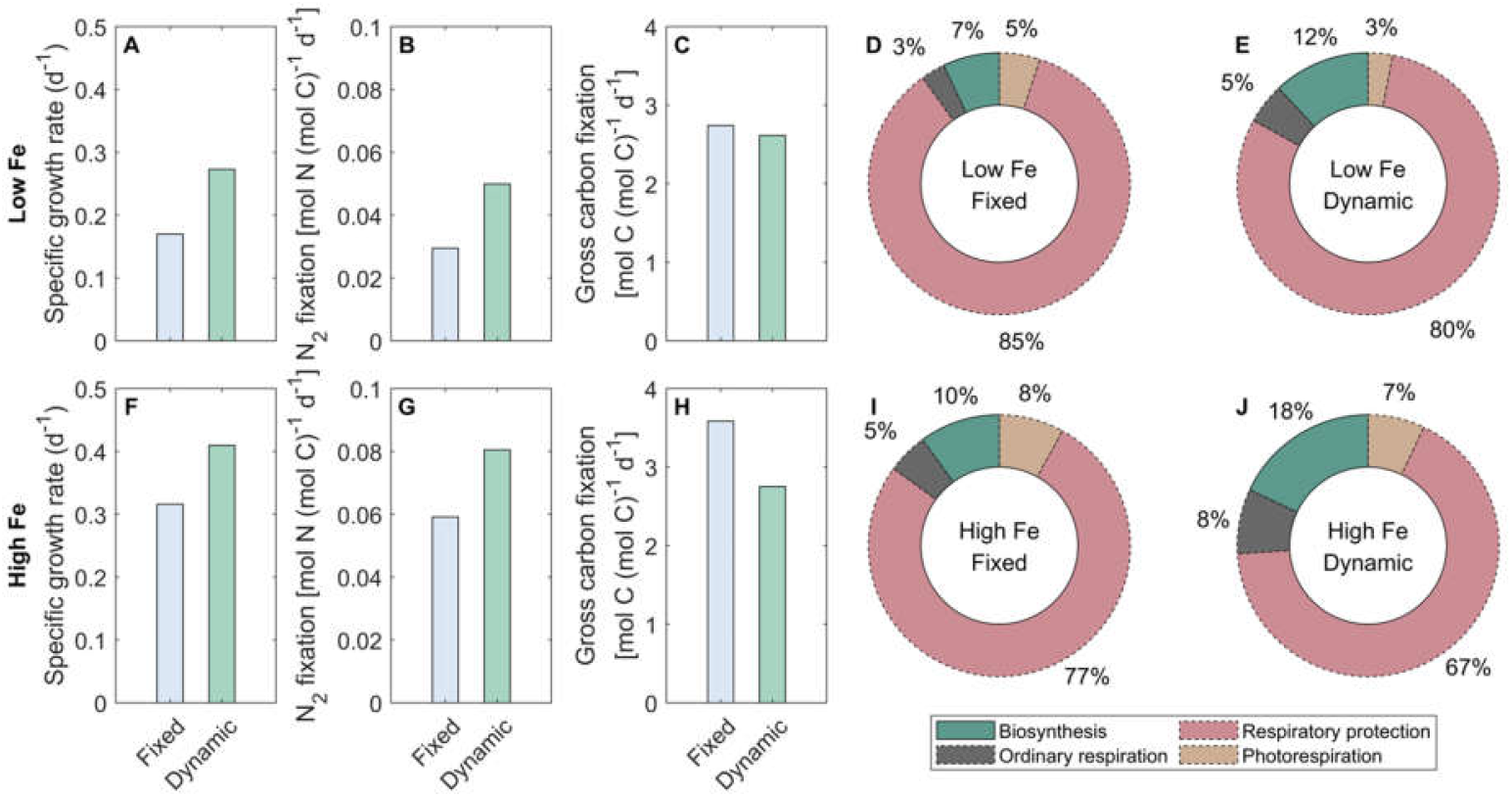
Modeled daily-integrated results of *Trichodesmium*. The model is simulated with diurnally fixed and dynamic O_2_ permeability of the cell membrane under low-Fe (40 pM) and high-Fe (1250 pM) conditions. The number in the inner circle represents the daily-integrated gross carbon fixation rate [mol C (mol C)^-1^ d^-1^]. The fixed carbon is allocated to photorespiration, respiratory protection, ordinary respiration and biosynthesis. Carbon use efficiency: the fraction of gross fixed carbon allocated to biosynthesis, which is highlighted using solid lines (**D, E, I** and **J**).

DPO_2_ benefits modeled *Trichodesmium* via modulating carbon and N_2_ fixation rates and intracellular O_2_ levels, with the physiological roles of DPO_2_ differs in two periods (Fig. 3). We first analyzed results under low-Fe conditions.

**FIG 3.**
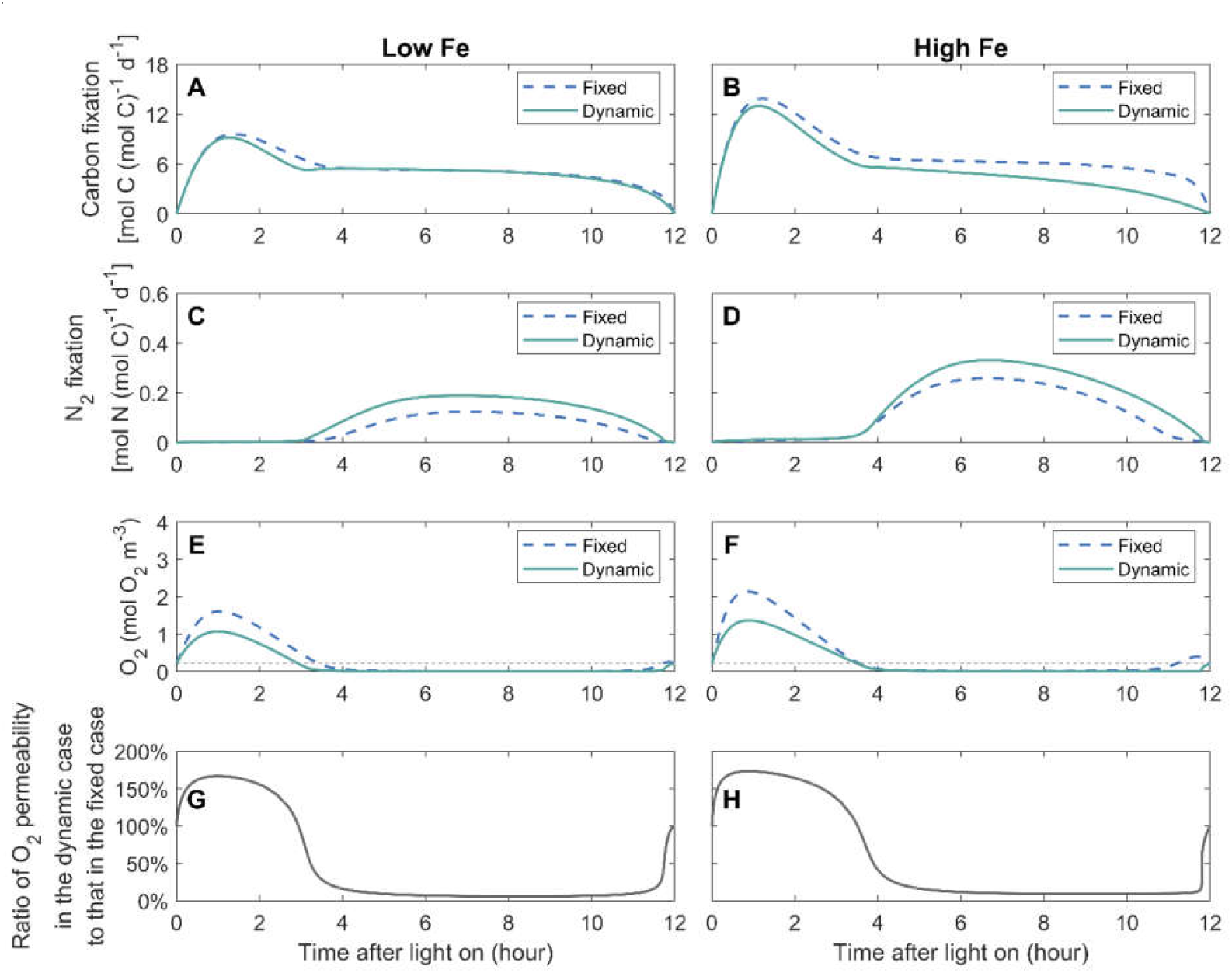
Simulated instantaneous rates of gross carbon fixation and N_2_ fixation, intracellular O_2_ concentrations, and O_2_ permeability of cell membrane during the light period. The O_2_ permeability is shown as the ratio of its values in the dynamic-permeability case to those in the fixed-permeability case. The model is simulated with diurnally fixed and dynamic O_2_ permeability of the cell membrane under low-Fe (40 pM) (**A, C, E** and **G**) and high-Fe (1250 pM) (**B, D, F** and **H**) conditions. The thin black dashed lines represent the ambient far-field O_2_ concentration.

During the early period, comparing to the fixed-permeability case, the carbon fixation rate in the dynamic-Fe case was slightly lower with a decreasing pattern (approximately 1−4 h) (Fig. 3A). This indicates that DPO_2_ could lower the requirement for carbon storage. Additionally, lower intracellular O_2_ level in the dynamic-permeability case (Fig. 3E, G) reduced the carbon consumption by photorespiration (*see* Discussion).

During the low-O_2_ window, the N_2_ fixation rate in the dynamic-permeability case was higher (Fig. 3C). Wider low-O_2_ window was presented in the dynamic-permeability case (Fig. 3E), suggesting DPO_2_ could reduce the stress from O_2_ on N_2_ fixation (*see* Discussion).

Under the high-Fe condition, diurnal patterns of modeled carbon and N_2_ fixation rates and intracellular concentrations were similar to those under the low-Fe condition, but at slightly higher levels (Fig. 3).

### Simulated diurnal intracellular O_2_ fluxes

The physiological functions of DPO_2_ in regulating intracellular O_2_ fluxes varied diurnally. During the early daytime (approximately 0−3 h of the light period), the daily-integrated net O_2_ production rate by PET was slightly lower compared to the dynamic-permeability case under low Fe (Fig. 4A), with more pronounced difference under high Fe (Fig. 4B) (*see* Discussion). Intracellular O_2_ diffused out of the cell cytoplasm (Fig. 4C, D) due to the high intracellular O_2_ concentration (Fig. 3E, F), induced by the high net O_2_ production rate of PET (Fig. 4A, B). The intracellular O_2_ concentration in the dynamic-permeability case was lower than that in the fixed-permeability case (Fig. 3E, F), while the physical diffusion rates of O_2_ in both cases were similar (Fig. 3C, D). This can be attributed to the higher O_2_ permeability of the cell membrane in the dynamic-permeability case (Fig. 3G, H), which facilitated quick intracellular O_2_ diffusion into extracellular environment.

**FIG 4.**
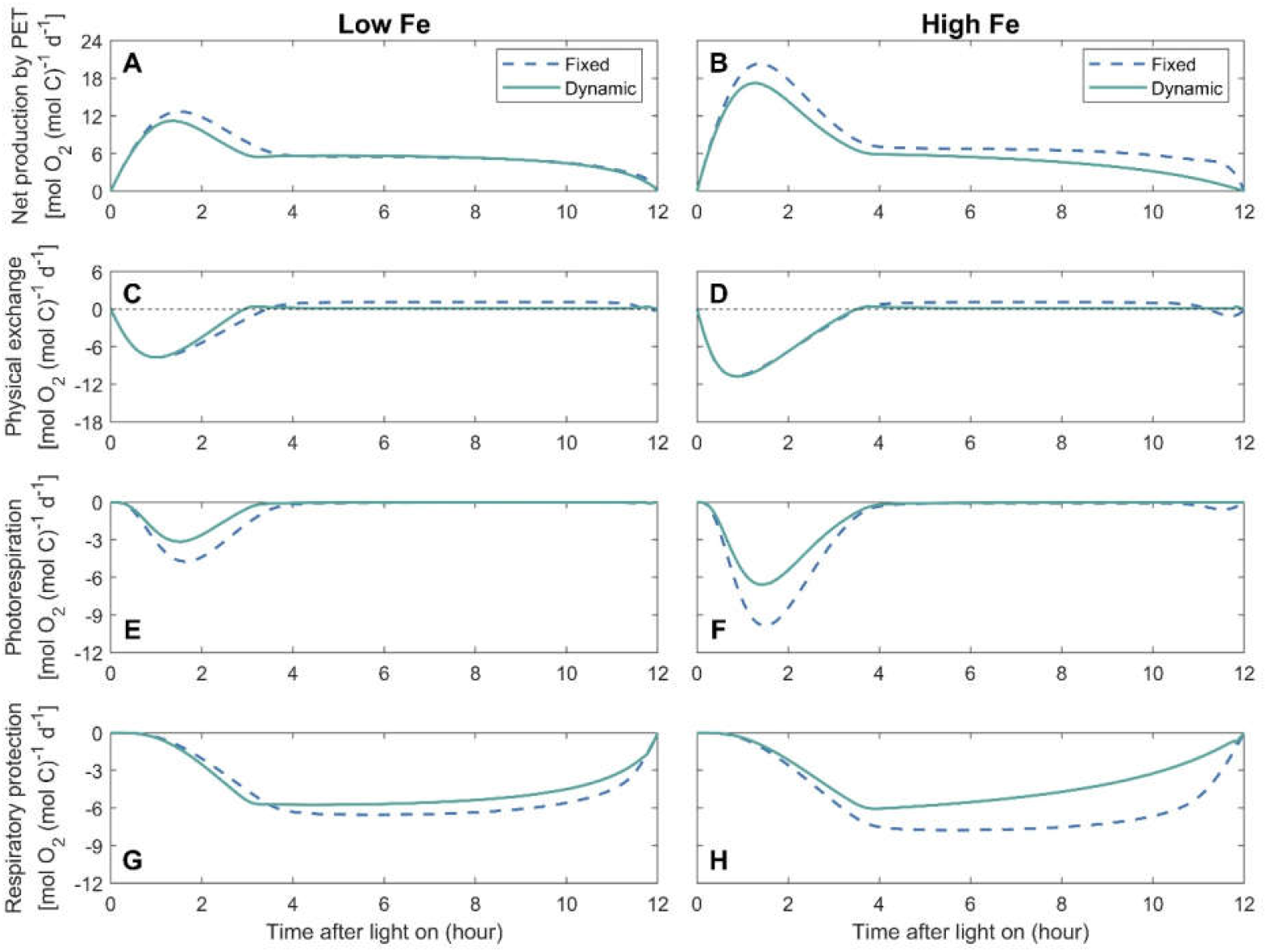
Simulated diurnal intracellular O_2_ fluxes. O_2_ fluxes include net production by photosynthetic electron transfer (PET), physical exchange between intracellular and extracellular environments, O_2_ consumption by photorespiration and respiratory protection (negative values). Positive physical O_2_ exchange represents O_2_ flux into cells. The model is simulated with diurnally fixed or dynamic O_2_ permeability of the cell membrane under low-Fe (40 pM) (**A, C, E** and **G**) and high-Fe (1250 pM) (**B, D, F** and **H**) conditions.

In addition, high intracellular O_2_ concentrations during this period stimulated photorespiration (Fig. 4E, F). The lower photorespiration rates in the dynamic-permeability case (Fig. 4E, F) were attributed to reduced intracellular O_2_ concentration (Fig. 3E, F). Therefore, DPO_2_ has the potential to alleviate the stress caused by photorespiration, particularly during the early daytime when net O_2_ production rate was high.

In the following light period, the intracellular low-O_2_ window was created (Fig. 3E, F) due to the downregulation of the net O_2_ production by PET (Fig. 4A, B) and the high O_2_ consumption by respiratory protection (Fig. 4G, H) (10), allowing the occurrence of N_2_ fixation. Extracellular O_2_ diffused into the cytoplasm (Fig. 4C, D), which was slowed down by the lowered O_2_ permeability in the dynamic-permeability case (Fig. 3G, H). Therefore, DPO_2_ saved organic carbon required by respiratory protection (Fig. 4G, H) to maintain low O_2_ levels in *Trichodesmium* (Fig. 3). In both the dynamic- and fixed-permeability cases, the photorespiration rate approached zero during this period (Fig. 4E, F), indicating that the creation of the low-O_2_ window also helped to mitigate O_2_-induced stresses on photosynthesis.

## DISCUSSION

In this study, we developed a physiological model of *Trichodesmium* trichome to quantitatively investigate the impact of photorespiration and the physiological advantages of DPO_2_ (Fig. 1). Intracellular O_2_ management such as respiratory protection was considered, with the temporal segregation between photosynthesis and N_2_ fixation formed and the low-O_2_ window created (Fig. 3). These were in line with previous model studies and observations (4, 9, 10, 12, 20). The model results demonstrate that DPO_2_ enhanced *Trichodesmium* N_2_ fixation and growth rates (Figs. 2, 3, 5 and S3) by decreasing photorespiration and respiratory protection and increasing carbon and Fe use efficiency.

**FIG 5.**
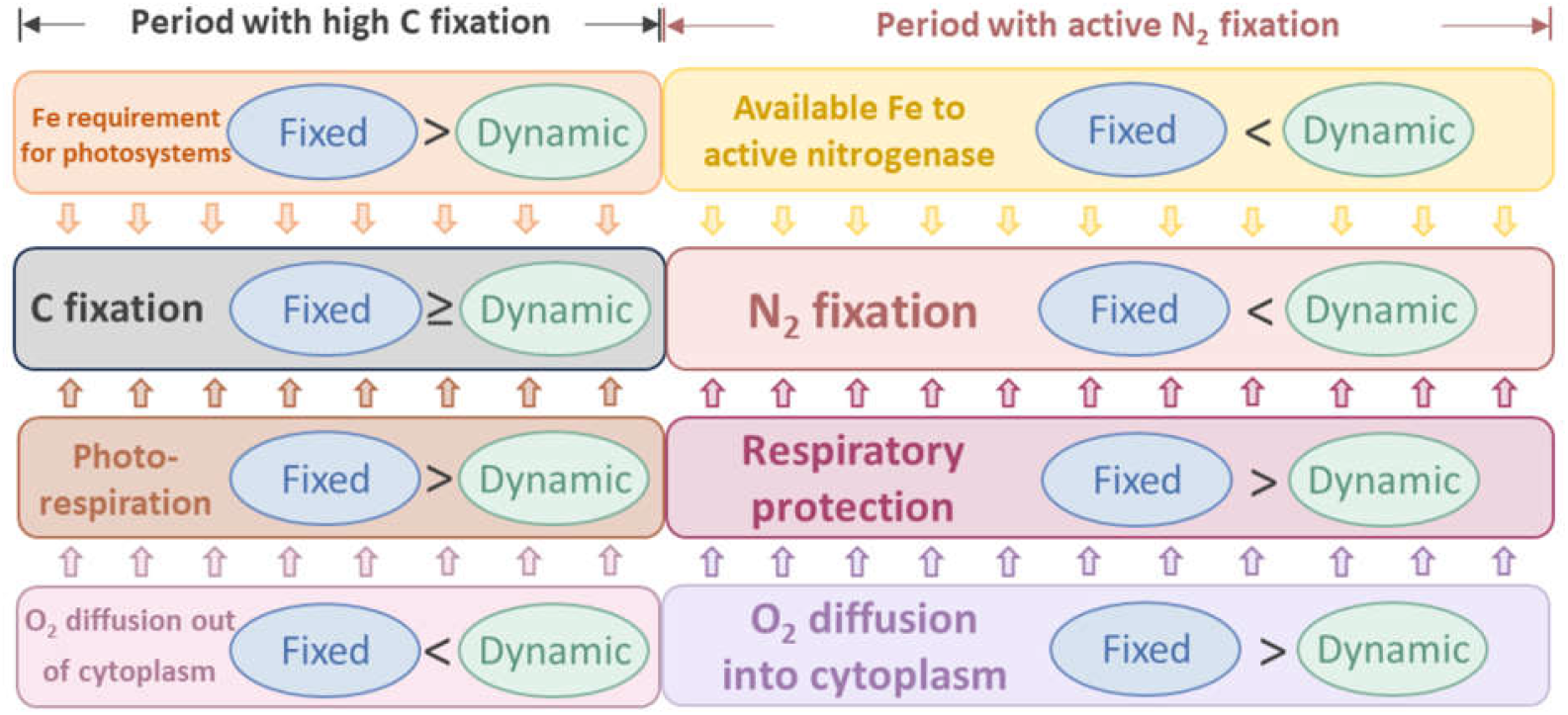
Schematic diagram illustrating diurnally fixed and dynamic O_2_ permeability of the cell membrane. Dynamic O_2_ permeability can reduce photorespiration and the requirement for respiratory protection. In addition, it can promote carbon and iron use efficiency. Arrows between boxes represent the influence toward carbon fixation and N_2_ fixation.

### Dynamic O_2_ permeability lowers photorespiration

In the model case when the O_2_ permeability was fixed, 5% and 8% of the daily-integrated gross fixed carbon were consumed by photorespiration under low-Fe and high-Fe conditions, respectively (Figs. 2 and 4E, F), which is comparable to the fractions allocated to biomass synthesis (Fig. 2). These low proportions of photorespiration seem to be reasonable in *Trichodesmium*, which operate carbon concentrating mechanisms (40) to enhance carbon fixation and minimize the oxygenation reaction (i.e., photorespiration) by RuBisCO (14). After implementing DPO_2_, modeled photorespiration rates were reduced by 42% and 35% in the dynamic-permeability case, respectively (Figs. 2, 4E, F and 5), resulting in a corresponding increase in gross fixed carbon allocated to biomass synthesis (Fig. 2). This was also found in the model experiments under constant light intensity (Fig. S4B). Further model experiments demonstrated that substituting the photorespiration rates in the dynamic-permeability case with those derived from the fixed-permeability case led to a reduction in *Trichodesmium* growth rates by 10% and 9% under low- and high-Fe conditions, respectively. These findings indicate that DPO_2_ has the potential to improve carbon use efficiency and growth rate of *Trichodesmium* by reducing its photorespiration.

The reduction of photorespiration in the dynamic-permeability case (Fig. 4E, F) was partially attributed to the increased O_2_ permeability during the early daytime (Figs. 3G, H and 5), facilitating the diffusion of intracellular O_2_ into the extracellular environment and resulting in lower intracellular O_2_ concentration (Fig. 3E, F).

In addition, DPO_2_ also resulted in decrease in photosynthesis and O_2_ production evolution during the early light period (Figs. 3A, B and 4A, B), contributing to reducing the intracellular O_2_ concentration (Fig. 3E, F) and photorespiration rate (Figs. 4E, F and 5). This can be attributed to the reduced need for respiratory protection during the low-O_2_ window (Figs. 4G, H and 5), thereby downregulating the requirement for carbon fixation (Figs. 3A, B and 5) and Fe allocated to photosystems (Fig. S5A, B) during the early daytime (discussed later).

Laboratory experiments on *Trichodesmium* have shown that light-dependent O_2_ uptake (which involves photorespiration as well as Mehler reaction and potentially flv-mediated O_2_ uptake) can indeed be dynamically regulated by external O_2_ and CO_2_ concentrations: When external O_2_ levels in the media were decreased or increased, light-dependent O_2_ uptake changed proportionally within minutes in a reversible fashion (9). Another recent culture study of *Trichodesmium* also proposed a decrease in photorespiration under low-O_2_ conditions (< 0.213 mol O_2_ m^-3^), although it was not quantified (16). Additionally, the rate of light-dependent O_2_ consumption has been shown to increase with an increase of CO_2_ concentration in the medium (41)

Another previous study proposed that photorespiration could serve as a mechanism to protect nitrogenase and N_2_ fixation by consuming the O_2_ produced by LPET, which, however, lacked support from observations or model simulations (42). In our model study, photorespiration primarily occurred during the early light period (Fig. 4E, F), when it was stimulated by high intracellular O_2_ concentrations (Fig. 3E, F), while there was minimal photorespiration during the low-O_2_ window when N_2_ fixation predominantly occurred (Fig. 4E, F). This low photorespiration could occur when high levels of respiratory protection depleted O_2_ and produced CO_2_ in *Trichodesmium* (Fig. 4G, H) (14), although the intracellular CO_2_ concentration was not simulated in our study. In other words, this suggests that respiratory protection played a major role lowering the intracellular O_2_ level for N_2_ fixation, while photorespiration may only have a limited contribution.

### Dynamic O_2_ permeability reduces the requirement for respiratory protection

In the fixed-permeability case, 85% and 77% of the gross fixed carbon were allocated to respiratory protection to consume intracellular O_2_ and create the low-O_2_ window for N_2_ fixation under low-Fe and high-Fe conditions, respectively (Figs. 2 and 4G, H). These percentages were generally consistent with previous model studies that adopted fixed O_2_ permeability of the cell membrane over the light period (10, 12, 20) or even other N_2_ fixers (e.g., 43, 44). In comparison, in the dynamic-permeability case, the fractions of gross fixed carbon allocated to respiratory protection were reduced to 80% and 67% under low-Fe and high-Fe conditions, respectively (Figs. 2 and 4G, H). This reduction in respiratory protection was primarily attributed to the low-O_2_ permeability in the dynamic-permeability case, which decreased the diffusion rate of extracellular O_2_ into cytoplasm (Figs. 4C, D and 5). As a result, the intracellular stress on N_2_ fixation caused by extracellular O_2_ was basically relieved. Consequently, lower rates of O_2_-consuming respiratory protection (Figs. 4G, H and 5) were required to create and maintain lower intracellular O_2_ concentrations (Figs. 3E, F and S4A, B), thereby supporting higher N_2_ fixation rates in the dynamic-permeability case (Figs. 3A, B and 5).

Previous model studies have proposed that respiratory protection is a crucial strategy for managing the intracellular O_2_ level and creating the low-O_2_ condition for N_2_ fixation in *Trichodesmium* and other N_2_-fixing cyanobacteria, such as *Crocosphaera* (4, 10, 44-46). Consistent with the previous study, our study demonstrated that respiratory protection respired major gross fixed carbon to consume intracellular O_2_, resulting in a high indirect cost for N_2_ fixation. This is also supported by observations of high daily-integrated gross fixed C:N ratios (e.g., 30–50), even when Fe is replete (4, 9, 47, 48). Lowering carbon demand may allow this high ratio of carbon to be channeled into growth, thus increasing ecological competitiveness. Therefore, *Trichodesmium* might develop several strategies to reduce the requirement for respiratory protection and promote the carbon use efficiency, such as the dynamic Fe allocation, of which the function was quantified in recent model study (20). On top of that, our study highlights the role of DPO_2_ in lowering respiratory protection and alleviating the stress from O_2_ on N_2_ fixation in *Trichodesmium*.

### Dynamic O_2_ permeability improves Fe use efficiency

The dynamic-permeability model case exhibited higher rates of N_2_ fixation and growth compared to the fixed-permeability case under both low and high Fe levels (Figs. 2, 3, 5 and S3), thereby promoting Fe use efficiency by 69% and 36%, respectively (Fig. 6).

**FIG 6.**
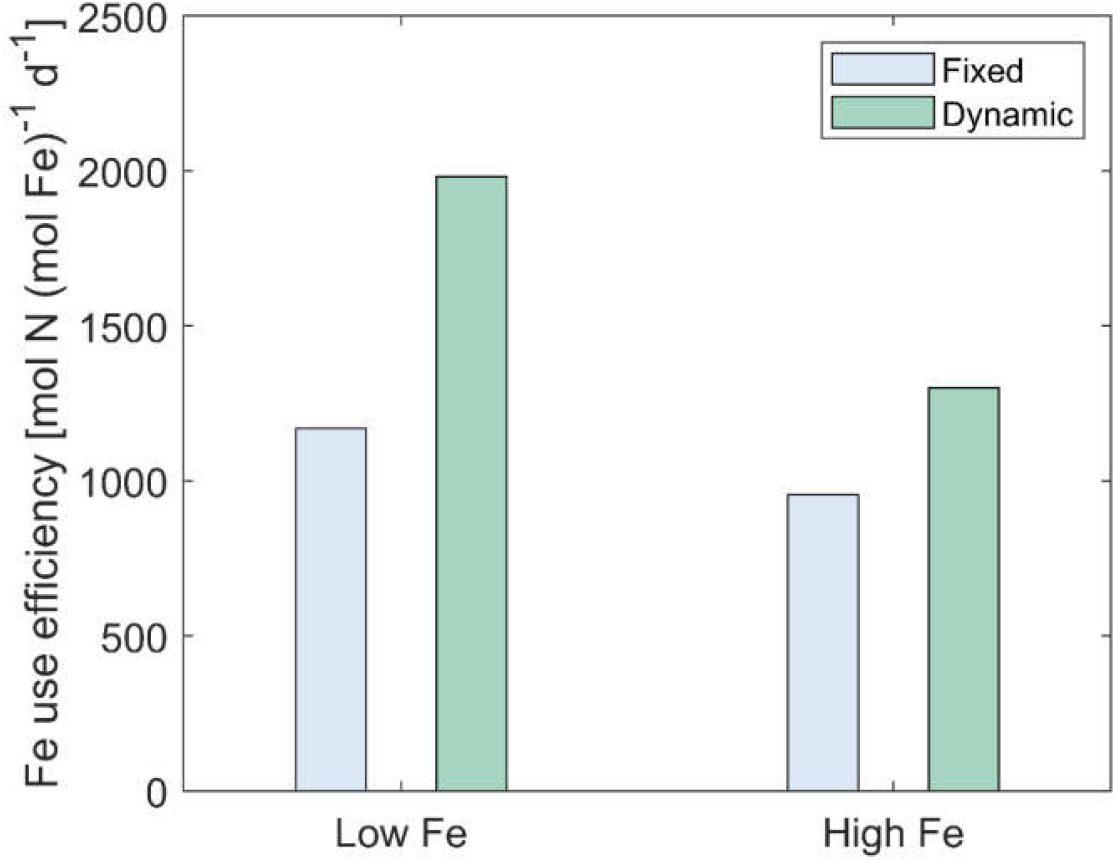
Modeled results of Fe use efficiency in *Trichodesmium*. The model is simulated with diurnally fixed and dynamic O_2_ permeability of the cell membrane under low-Fe (40 pM) and high-Fe (1250 pM) conditions. Fe use efficiency: the ratio of daily-integrated N_2_ fixation rate to intracellular metabolic Fe.

The simulated diurnal patterns of Fe in both photosystems and nitrogenase suggest that DPO_2_ could regulate intracellular Fe allocation (Fig. S5). Specifically, in the dynamic-permeability case, less Fe was allocated to photosystems during the light period compared to the fixed-permeability case (Fig. S5A, B). This can be attributed to two main reasons. First, the reduction in the photorespiration allowed for the saving of energy and NADPH, which then can be utilized by carbon fixation. Second, the downregulation of respiratory protection decreased the demand for carbon fixation and storage in early daytime (Figs. 3 and 5). In addition, decreased respiratory protection mitigated the inhibition on photosynthetic electron transfer (4). Consequently, the Fe demand of photosystems was reduced in the dynamic-permeability case (Figs. 5 and S5A, B). The saved Fe from photosystems could then be allocated to (active) nitrogenase (Figs. 5 and S5C–F), enhancing N_2_ fixation (Fig. 3C, D and 5).

In conclusion, our model study highlights two main potential mechanisms explaining the benefits of dynamic O_2_ permeability to N_2_ fixation and growth in *Trichodesmium*, including the reduced photorespiration and decreased requirement for respiratory protection (Fig. 5). This can improve carbon and Fe use efficiency (Figs. 5 and 6), as well as N_2_ fixation and growth rates of *Trichodesmium*, especially under low Fe. Therefore, this strategy potentially alleviates Fe limitation, helping *Trichodesmium* survive in the oligotrophic open ocean.

It should be noted that in our model, variations in the O_2_ permeability of *Trichodesmium* cell membrane were assumed to occur instantly in response to intracellular O_2_ levels. However, further investigation is required to explore the timescale and the efficiency of redistributing hopanoid lipids in the cell membrane of *Trichodesmium*, which regulates the O_2_ permeability. This will provide a more comprehensive understanding of the physiological benefits associated with DPO_2_ in *Trichodesmium*.

### Broader context: Is dynamic O_2_ permeability a common strategy in marine cyanobacterial diazotrophs?

The gene for the synthesis of hopanoids (the squalene-hopene cyclase gene, *shc*) was reported to be present in UCYN-A (nitroplast), UCYN-B (*Crocosphaera*) and UCYN-C (*Cyanothece*) (18, 49-54), suggesting their potentials in dynamically regulating the membrane permeability to O_2_ (Table 1). Hopanoids are known to decrease membrane O_2_ permeability, providing protection to intracellular processes such as nitrogenase activity (18). However, direct evidence of diel variation in hopanoid production or dynamic changes in membrane lipid composition remains limited in marine diazotrophs.

**TABLE 1.** Summary of indications from literature for dynamic O_2_ permeability with potential benefits in some marine cyanobacterial diazotrophs.

| Diazotroph | Indications/Potential for<br>dynamic O<sub>2</sub> permeability | References |
| --- | --- | --- |
| *Trichodesmium* | Yes | This study and (18) |
| UCYN-A (nitroplast) | Possible | (51) |
| UCYN-B (*Crocospaera*) | Yes | (49) |
| UCYN-C (*Cyanothece*) | Yes | (49) |
| Diazotroph-diatom association (DDA) | No | (18) |

A recent transcriptomic study (55) reveals diurnal cycling of gene expression in *Crocosphaera*, including genes related to photosynthesis and N_2_ fixation, suggesting that membrane lipid composition may be dynamically regulated over diel cycles. Similarly, in non-diazotrophic cyanobacteria like *Synechocystis* sp. PCC 6803, diel light-dark cycles significantly influence fatty acid synthesis and membrane lipid turnover (56). Beyond cyanobacteria, studies in plants show that membrane lipid turnover can occur on timescales as short as two hours (57), suggesting that dynamic regulation of membrane lipids is a widespread strategy across diverse organisms. Together, these findings support the plausibility of DPO_2_ as a physiological strategy in marine diazotrophs.

While the diel rhythms of photosynthesis and N_2_ fixation in these marine non-heterocyst-forming diazotrophic cyanobacteria may differ from *Trichodesmium* (58, 59), it is possible that the proposed physiological advantages of DPO_2_ may apply to these diazotrophs.

The physiological roles of DPO_2_ in improving the carbon and Fe use efficiency in some marine diazotrophs were similar to those of the dynamic Fe allocation (20). This suggests that dynamic regulation of membrane permeability can facilitate N_2_ fixation and the growth of these diazotrophs in a manner similar to dynamic regulation of physiological processes, including temporal segregation (58, 60) and potential rapid mode switching (61) between photosynthesis and N_2_ fixation. The in-depth controlling mechanisms of dynamic strategies guarantees further research.

#### UCYN-A

The genome of UCYN-A (or nitroplast) (53) is highly reduced and lacks genes for O_2_-producing photosystem II (62), suggesting a potential reduction in O_2_-induced stresses and photorespiration. A previous study demonstrated coordination between the expression of the *shc* gene and the *nifH* gene (encoding nitrogenase) in UCYN-A (49). This indicates that UCYN-A may regulate membrane permeability to O_2_ dynamically, lowering intracellular O_2_ levels and protecting nitrogenase activity by mitigating O_2_ diffusion from the O_2_-producing haptophyte algal host cell and the ambient environment into the cytoplasm during active N_2_ fixation. In addition, the low transcription level of cytochrome *c* oxidase *coxA* gene in UCYN-A during the light period (24, 49) indicates a reduced requirement for respiratory protection. According to insights from this study, such dynamic regulation could enhance carbon and Fe use efficiency in UCYN-A.

#### UCYN-B and UCYN-C

Both UCYN-B (*Crocosphaera*) and UCYN-C (*Cyanothece*) conduct photosynthesis during the light period and N_2_ fixation at night (49, 63-65). Transcriptomic analysis has shown that the *shc* gene in UCYN-B and UCYN-C reached its peak expression just before the increase in the *nifH* gene expression level (49). This observation suggests that UCYN-B and UCYN-C may produce hopanoids to protect N_2_ fixation from O_2_ diffusing from the external environment during the dark period, therefore reducing the level of respiratory protection required to safeguard nitrogenase and support N_2_ fixation (44, 45). Additionally, lower expression of the *shc* gene during the daytime in UCYN-B and UCYN-C (49) might result in higher O_2_ permeability, facilitating the diffusion of photosynthetically produced O_2_ out of the cell and thus reducing photorespiration. These mechanisms, as proposed in this study, would lead to elevated carbon and Fe use efficiency. Additionally, these organisms have heterogeneous rates of N_2_ fixation (i.e., some cells fix N_2_ and others do not) (66), and thus, the most effective DPO_2_ would also be heterogeneous across their population. Furthermore, the presence of hopanoid rafts in UCYN-B (67) suggests that hopanoid may be redistributed within the membrane of to dynamically modulate O_2_ permeability (68).

#### DDAs

Heterocyst-forming diazotrophs (diatom-diazotroph assemblage, DDA) lack the *shc* gene responsible for hopanoid synthesis (18). This indicates that DPO_2_ may not be necessary in DDA. Instead, the heterocyst use glycolipids to form barrier against extracellular O_2_ (69, 70), seemingly providing sufficient protection for nitrogenase activity and reducing Fe requirements compared to non-heterocyst-forming cyanobacterial diazotrophs (71). Similarly, the absence of the *shc* gene in non-diazotrophic cyanobacteria such as *Prochlorococcus* and *Synechococcus* (18) further supports the idea that DPO_2_ is one of the evolved strategies of diazotrophs when facing O_2_ stress on N_2_ fixation.

## Conclusions

We investigated how dynamic cell permeability to O_2_ (DPO_2_) in *Trichodesmium* trichomes may enhance N_2_ fixation and growth rates, taking into account the effect of photorespiration. Our model shows that DPO_2_ reduces photorespiration especially during the early daytime, lowering the requirement for respiratory protection, facilitating the formation of the low-O_2_ intracellular condition for N_2_ fixation and improving carbon use efficiency. Moreover, DPO_2_ in *Trichodesmium* may impact the diurnal allocation of the intracellular Fe, ultimately promoting the Fe use efficiency. Fragmental evidences for DPO_2_ are reported in other marine diazotrophs, suggesting that DPO_2_ is a common strategy adopted by marine diazotrophs. The model framework presented in our study could also be used to explore other physiological mechanisms that control N_2_ fixation, such as light and Fe colimitation. It can also be incorporated to biogeochemical models to enhance their predictive capabilities in the ecophysiology of marine diazotrophs.

## Supporting information

SUPPLEMENTARY METHODS: Full model description. Table S1-S3. FIG S1-S5

## DATA AND MATERIALS AVAILABILITY

All data, code and materials used in this study are available from the corresponding author upon reasonable request. The code is freely available in Zenodo once the manuscript is published.

## AUTHOR CONTRIBUTIONS

WL and YWL originated concept for the study. WL designed numerical model. WL coded the initial version of the model and performed numerical modeling. WL, KI, OP, ME, and YWL analyzed results and improved the numerical model. WL wrote the first draft of the manuscript, and all coauthors revised the manuscript.

## DECLARATION OF COMPETING INTEREST

The authors declare that they have no competing interests.

## ACKNOWLEDGMENTS

This project is partly supported by the National Natural Science Foundation of China (42076153 and 42376140 to YWL), China Scholarship Council, and MEL PhD Fellowship to WL. This work was supported by a grant from the Simons Foundation (LS-ECIAMEE-00001549, Inomura). OP is supported by GACR 23-06593S and by OP JAK Photomachines. ME is supported by GACR GA24-12396S and by OP JAK project Photomachines.

## Notes

### Competing Interest Statement

The authors have declared no competing interest.

