## SUPPLEMENTARY METHODS: Full model description. Table S1-S3. FIG S1-S5 for "Modeling dynamic oxygen permeability as a mechanism to mitigate oxygen-induced stresses on photosynthesis and N_2_ fixation in marine *Trichodesmium*"

#### 1. Photosynthetic pathways

Linear photosynthetic electron transfer (LPET) and alternative electron transfer (AET) are simulated in our model (Fig. 1A). Because (Mehler reaction)-mediated AET is the dominant AET in *Trichodesmium* (1), other AET (e.g., the cyclic electron transfer around photosystem I, the AET mediated by midstream oxidase with photosystem II, and the AET from photosystem II to the respiratory terminal oxidase respiratory terminal oxidase) (2), are not considered.

Both LPET and AET generate a proton gradient of 12 $H^+$ per 8 photons (3). By assuming that 3 ATP are produced by the thylakoid ATP synthase with a proton gradient of 14 $H^+$, for 4 electrons through PET, LPET produces 2.6 ATP, with 2 NADPH and 1 $O_2$, while AET only produces 2.6 ATP (2-4).

The total PET rate [$V_{PET}$, mol electron (mol C)$^{-1}$ s$^{-1}$] is regulated by light intensity ($I$, µmol m$^{-2}$ s$^{-1}$) and the Fe quota in photosystems [$Fe_{PS}$, µmol Fe (mol C)$^{-1}$], and $V_{PET}$ is inhibited by respiratory protection (RP) [$V_{RP}$, mol C (mol C)$^{-1}$ s$^{-1}$, described later] (5).

$$V_{PET} = v_{PET}^{max} \cdot \frac{Fe_{PS}}{Fe_{PS} + k_{Fe}^{PS}} \cdot (1 - e^{-\alpha_I \cdot I}) \cdot e^{-\beta \cdot V_{RP}}, \quad \text{(S1)}$$

where $v_{PET}^{max}$ [mol electron (mol C)$^{-1}$ s$^{-1}$] is the maximal rate of PET, $k_{Fe}^{PS}$ [µmol Fe (mol C)$^{-1}$] is the half-saturating coefficient of $Fe_{PS}$ for PET, $\alpha_I$ (µmol$^{-1}$ m$^2$ s) is the initial slope of PET versus light curve, and $\beta$ [mol C (mol C)$^{-1}$ s] represents the degree of the inhibition effect from RP on PET. Note that in fixed-Fe and dynamic-Fe cases, the light intensity is diurnally variant with time, following the sine function during a 12-hour light period in our model (6).

To determine the rates of LPET and AET [$V_{LPET}$ and $V_{AET}$ , mol electron (mol C)$^{-1}$ s$^{-1}$], the fraction of photosynthetic electrons flowing into AET ($f_{AET}$, dimensionless) is introduced in our model, and $f_{AET}$ is calculated in each time step to fulfill the intracellular immediate requirement of ATP and NADPH (7).

$$V_{LPET} = V_{PET} \cdot (1 - f_{AET}), \quad \text{(S2)}$$

$$V_{AET} = V_{PET} \cdot f_{AET}. \quad \text{(S3)}$$

The NADPH production rate [$V_{NAPDH}$, mol NADPH (mol C)$^{-1}$ s$^{-1}$] is:

$$V_{NAPDH} = V_{LPET} \cdot q_{LPET}^{NADPH}, \quad \text{(S4)}$$

where $q_{LPET}^{NADPH}$ = 0.5 mol NADPH (mol electron)$^{-1}$ is the ratio of NADPH to electron in LPET (4).

The ATP production rate [$V_{ATP}$, mol ATP (mol C)$^{-1}$ s$^{-1}$] is:

$$V_{ATP} = V_{LPET} \cdot q_{LPET}^{ATP} + V_{AET} \cdot q_{AET}^{ATP}, \quad \text{(S5)}$$

where $q^{ATP}_{LPET} = q^{ATP}_{AET} = 0.65$ mol ATP (mol electron)$^{-1}$ ratios of ATP to electron in LPET and AET (2-4).

The $O_2$ production rate [$V_{O_2}$, mol $O_2$ (mol C)$^{-1}$ s$^{-1}$] is:

$$V_{O_2} = V_{LPET} \cdot q^{O_2}_{LPET}, \quad \text{(S6)}$$

where $q^{O_2}_{LPET} = 0.25$ mol $O_2$ (mol electron)$^{-1}$ is the ratio of $O_2$ to electron in LPET (3, 4).

### 2. Photorespiration

Photorespiration is a light-dependent process, which consumes ATP (adenosine triphosphate), NADPH (nicotinamide adenine dinucleotide phosphate hydrogen), $O_2$ and organic carbon, and produces $CO_2$ (8).

The maximal photorespiration rate [$V^{max}_{PR}$, mol C (mol C)$^{-1}$ s$^{-1}$] is computed based on the consumption that produced ATP by PET is fully consumed by photorespiration:

$$V^{max}_{PR} = \frac{V_{ATP}}{q^{ATP}_{PR}}, \quad \text{(S7)}$$

where $q^{ATP}_{PR} = 7$ mol ATP (mol C)$^{-1}$ is ATP to C ratio in photorespiration (8).

The rate of photorespiration [$V_{PR}$, mol C (mol C)$^{-1}$ s$^{-1}$] is also regulated by intracellular carbohydrate [$CH_2O$, mol C (mol C)$^{-1}$] and $O_2$ [$O_2$, mol $O_2$ m$^{-3}$]:

$$V_{PR} = V^{max}_{PR} \cdot \frac{CH_2O}{CH_2O + k^{PR}_{CH_2O}} \cdot \frac{O_2}{O_2 + k^{PR}_{O_2}}, \quad \text{(S8)}$$

where $k^{PR}_{CH_2O} = 0.4$ [mol C (mol C)$^{-1}$] and $k^{PR}_{O_2} = 1.92$ (mol $O_2$ m$^{-3}$) are half-saturating coefficients of $CH_2O$ and $O_2$ for photorespiration (Fig. S1B).

The NADPH, ATP and $O_2$ consumption rates of photorespiration [$V^{PR}_{NADPH}$, $V^{PR}_{ATP}$ and $V^{PR}_{O_2}$, mol NADPH (mol C)$^{-1}$ s$^{-1}$, mol ATP (mol C)$^{-1}$ s$^{-1}$ and mol $O_2$ (mol C)$^{-1}$ s$^{-1}$] are:

$$V^{PR}_{NADPH} = V_{PR} \cdot q^{NADPH}_{PR}, \quad \text{(S9)}$$

$$V^{PR}_{ATP} = V_{PR} \cdot q^{ATP}_{PR}, \quad \text{(S10)}$$

$$V^{PR}_{O_2} = V_{PR} \cdot q^{O_2}_{C,PR}, \quad \text{(S11)}$$

where $q^{NADPH}_{PR} = 4$ mol NADPH (mol C)$^{-1}$ and $q^{O_2}_{C,PR} = 3$ mol $O_2$ (mol C)$^{-1}$ are NADPH to C and $O_2$ to C ratios in photorespiration (8).

### 3. $N_2$ fixation

$N_2$ fixation requires both ATP and NADPH (9, 10). The maximal $N_2$ fixation rate [$V_{NF}^{max}$, mol N (mol C)$^{-1}$ s$^{-1}$] is calculated based on the assumption that produced ATP of PET are completely consumed by $N_2$ fixation.

$$V_{NF}^{max} = \frac{V_{ATP}}{q_{NF}^{ATP}}, \tag{S12}$$

where $q_{NF}^{ATP}$ = 9 mol ATP (mol N)$^{-1}$ is ATP:N ratio in $N_2$ fixation (9, 10).

The rate [$V_{NF}$, mol N (mol C)$^{-1}$ s$^{-1}$] is also regulated by the Fe quota in nitrogenase [$Fe_{NF}$, μmol Fe (mol C)$^{-1}$] and inhibited by intracellular $O_2$.

$$V_{NF} = V_{NF}^{max} \cdot \frac{Fe_{NF}}{Fe_{NF} + k_{Fe}^{NF}} \cdot (1 - \frac{O_2}{O_2 + k_{O_2}^{NF}}), \tag{S13}$$

where $k_{Fe}^{NF}$ [μmol Fe (mol C)$^{-1}$] and $k_{O_2}^{NF}$ (mol $O_2$ m$^{-3}$) are half-saturating coefficients of $Fe_{NF}$ and $O_2$ for $N_2$ fixation.

The NADPH and ATP consumption rates of $N_2$ fixation [$V_{NADPH}^{NF}$ and $V_{ATP}^{NF}$, mol NADPH (mol C)$^{-1}$ s$^{-1}$ and mol ATP (mol C)$^{-1}$ s$^{-1}$] are:

$$V_{NADPH}^{NF} = V_{NF} \cdot q_{NF}^{NADPH}, \tag{S14}$$

$$V_{ATP}^{NF} = V_{NF} \cdot q_{NF}^{ATP}. \tag{S15}$$

where $q_{NF}^{NADPH}$ = 3 mol NADPH (mol N)$^{-1}$ is NADPH:N ratio in $N_2$ fixation (9, 10).

### 4. $CO_2$ concentrating mechanism and carbon fixation

The energy requirement of the $CO_2$ concentrating mechanism (CCM) [$q_{CCM}^{ATP}$ = 0.8 mol ATP (mol C)$^{-1}$] is calculated based on the fraction of leakage (50%) of inorganic carbon ($C_i$), the fraction of $HCO_3^-$ in $C_i$ (80%), and the cost for per $HCO_3^-$ transportation [0.5 mol ATP (mol C)$^{-1}$] (11, 12).

Energy consumption rate of CCM [$V_{ATP}^{CCM}$, mol ATP (mol C)$^{-1}$ s$^{-1}$] is:

$$V_{ATP}^{CCM} = V_{CF} \cdot q_{CCM}^{ATP}, \tag{S16}$$

where $V_{CF}$ [mol C (mol C)$^{-1}$ s$^{-1}$] is carbon fixation rate.

Carbon fixation also requires both NADPH and ATP (13), and consumption rates [$V_{NADPH}^{CF}$ and $V_{ATP}^{CF}$, mol NADPH (mol C)$^{-1}$ s$^{-1}$ and mol ATP (mol C)$^{-1}$ s$^{-1}$] are:

$$V_{NADPH}^{CF} = V_{CF} \cdot q_{CF}^{NADPH}, \tag{S17}$$

$$V_{ATP}^{CF} = V_{CF} \cdot q_{CF}^{ATP}. \tag{S18}$$

$V_{CF}$ is solved at each time step with $f_{AET}$, based on the consumption that total NADPH and ATP production by PET are immediately and fully utilized by intracellular process:

$$V_{NADPH} = V_{NADPH}^{PR} + V_{NADPH}^{CF} + V_{NADPH}^{NF}, \tag{S19}$$

$$V_{ATP} = (V_{ATP}^{CCM} + V_{ATP}^{PR} + V_{ATP}^{CF} + V_{ATP}^{NF}) \cdot (1 + \gamma_{MT}), \tag{S20}$$

where $\gamma_{MT}$ (dimensionless) represents the ratio of ATP consumption by maintenance to other processes.

The carbon skeleton production rate [$V_{CS}$, mol C (mol C)$^{-1}$ s$^{-1}$] is stimulated by carbohydrate and downregulated by its own accumulation [$CS$, mol C (mol C)$^{-1}$]:

$$V_{CS} = v_{CS}^{max} \cdot \frac{CH_2O}{CH_2O + k_{CH_2O}^{CS}} \cdot \frac{CS_{max} - CS}{CS_{max}}, \tag{S21}$$

where $v_{CS}^{max}$ [mol C (mol C)$^{-1}$ s$^{-1}$] is the maximal production rate of the carbon skeleton, $k_{CH_2O}^{CS}$ [mol C (mol C)$^{-1}$] is the half-saturation constant of carbohydrates for carbon skeleton production, and $CS_{max}$ [mol C (mol C)$^{-1}$] is the maximum CS storage.

### 5. Respiratory protection

Respiratory protection (RP) rate is regulated by the demand for $N_2$ fixation and intracellular $O_2$ (7, 14, 15):

$$V_{RP} = v_{RP}^{max} \cdot (1 - e^{-\alpha_I \cdot I}) \cdot \frac{CS}{CS + k_{CS}} \cdot \left(\frac{N_{max} - N}{N_{max}}\right) \cdot \frac{O_2}{O_2 + k_{O_2}^{NF}}, \tag{S22}$$

where $v_{RP}^{max}$ [mol C (mol C)$^{-1}$ s$^{-1}$] is the maximal respiratory protection rate, $k_{CS}$ [mol C (mol C)$^{-1}$] is the half-saturating coefficient of the carbon skeleton for respiratory protection, and $N_{max}$ [mol N (mol C)$^{-1}$] is the maximal N storage.

The $O_2$ consumption by RP [$V_{O_2}^{RP}$, mol $O_2$ (mol C)$^{-1}$ s$^{-1}$] is:

$$V_{O_2}^{RP} = V_{RP} \cdot q_C^{O_2}, \tag{S23}$$

where $q_C^{O_2}$ [mol $O_2$ (mol C)$^{-1}$] is the ratio of $O_2$ to carbon in carbohydrate respiration.

### 6. $O_2$ diffusion

The ratio ($\varepsilon$, dimensionless) of the $O_2$ diffusion coefficient of the cell membrane relative to that in seawater ($d_{O_2}$, m$^2$ s$^{-1}$) (i.e., parameter measuring the $O_2$ permeability of the membrane) is considered to change throughout the daytime (16). $\varepsilon$ is assumed to increase with the increase of intracellular $O_2$ concentration:

$$\varepsilon = \varepsilon_{max} \cdot \frac{O_2}{O_2 + k_{O_2}^{diff}}, \tag{S24}$$

where $\varepsilon_{max}$ is the maximal relative diffusion coefficient and $k_{O_2}^{diff}$ (mol $O_2$ m$^{-3}$) is the half-saturation constant of $O_2$ for $\varepsilon$.

The rate of $O_2$ diffusion ($T_{O_2}$, mol $O_2$ m$^{-3}$ s$^{-1}$) between intracellular cytoplasm and ambient environment is parameterized by adopting the scheme of (17):

$$T_{O_2} = \frac{-2\cdot\pi\cdot d_{O_2}\cdot L}{V} \cdot \left\{\frac{1}{\varepsilon}\cdot\ln\left(\frac{R}{R+L_g}\right) - \ln\left(\frac{R+L_g+L_b}{R+L_g}\right)\right\}^{-1} \cdot (O_2^E - O_2), \qquad \text{(S25)}$$

where $L$ (m) and $V$ (m$^3$) are the length and the volume of the trichome, $R$ (m) is the radius of the cytoplasm, $L_g$ (m) is the thickness of the cell membrane, $L_b$ (m) is the thickness of the boundary layer, $O_2^E$ is the ambient far-field $O_2$ concentration.

### 7. Intracellular Fe pools and translocation

*Trichodesmium* can uptake more Fe than that required by its metabolism (called 'luxury uptake') especially in high-Fe environments, and the excess Fe is stored for surviving in low Fe environments (18, 19). Therefore, the total intracellular Fe quota [$Fe$, µmol Fe (mol C)$^{-1}$] consists of Fe in metabolism and storage [$Fe_M$ and $Fe_{ST}$, µmol Fe (mol C)$^{-1}$], calculated based on the threshold of Fe [$Fe_{TH}$, µmol Fe (mol C)$^{-1}$] (20).

$$Fe_M = Fe, \qquad \text{when } Fe \le Fe_{TH}, \qquad \text{(S26)}$$

$$Fe_M = Fe_{TH} + (1 - f_{ST}) \cdot (Fe - Fe_{TH}), \text{ when } Fe > Fe_{TH}, \qquad \text{(S27)}$$

where $f_{ST}$ (dimensionless) is the fraction of luxury Fe uptake.

Fe allocations are among $Fe_M$, including Fe in photosystems, active nitrogenase, inactivated nitrogenase, maintenance and buffer [$Fe_{PS}$, $Fe_{NF}$, $Fe_{NF}^{NA}$, $Fe_{MT}$ and $Fe_{BF}$, µmol Fe (mol C)$^{-1}$] (Fig. 1B). Fe in maintenance is set diurnally constant at 10% of $Fe_M$ (20). Fe used in the photosystems and nitrogenase is from the buffer pool (15).

The synthesis rate of photosystems [$T_{PS}^{BF}$, µmol Fe (mol C)$^{-1}$ s$^{-1}$] is stimulated by light intensity and is gradually saturated with $Fe_{PS}$:

$$T_{PS}^{BF} = TF_{PS_{max}}^{BF} \cdot (1 - e^{-\alpha_I \cdot I}) \cdot (1 - \frac{Fe_{PS}}{Fe_{PS} + k_{Fe_{PS}}^{PS_{syn}}}), \qquad \text{(S28)}$$

where $T_{PS_{max}}^{BF}$ [µmol Fe (mol C)$^{-1}$ s$^{-1}$] is the maximal synthesis rate of photosystems, $k_{Fe_{PS}}^{PS_{syn}}$ [µmol Fe (mol C)$^{-1}$] is the half-saturating coefficients of $Fe_{PS}$ for the synthesis of photosystems.

The decomposition rate of photosystems [$T_{BF}^{PS}$, µmol Fe (mol C)$^{-1}$ s$^{-1}$] is stimulated by $Fe_{PS}$ but inhibited by respiratory protection (5):

$$T_{BF}^{PS} = T_{BF_{max}}^{PS} \cdot \frac{Fe_{PS}}{Fe_{PS} + k_{Fe_{PS}}^{PS_{dec}}} \cdot e^{-\beta \cdot V_{RP}}, \tag{S29}$$

where $T_{BF_{max}}^{PS}$ [μmol Fe (mol C)$^{-1}$ s$^{-1}$] is the maximal decomposition rate of photosystems, $k_{Fe_{PS}}^{PS_{dec}}$ [μmol Fe (mol C)$^{-1}$] is the half-saturating coefficient of $Fe_{PS}$ for the decomposition of photosystems. Fe released from decomposed photosystems returns to buffer pool (15).

For nitrogenase, intracellular requirement for $N_2$ fixation and $Fe_{BF}$ regulate its synthesis rate [$T_{NF}^{BF}$, μmol Fe (mol C)$^{-1}$ s$^{-1}$]:

$$T_{NF}^{BF} = T_{NF_{max}}^{BF} \cdot (1 - e^{-\alpha_I \cdot I}) \cdot \frac{CS}{CS + k_{CS}} \cdot \left(\frac{N_{max} - N}{N_{max}}\right) \cdot \frac{Fe_{BF}}{Fe_{BF} + k_{Fe_{BF}}^{NF_{syn}}}, \tag{S30}$$

where $T_{NF_{max}}^{BF}$ [μmol Fe (mol C)$^{-1}$ s$^{-1}$] is the maximal nitrogenase synthesis rate, $k_{CS}$ [mol C (mol C)$^{-1}$] and $k_{Fe_{BF}}^{NF_{syn}}$ [μmol Fe (mol C)$^{-1}$] are half-saturating coefficients of the carbon skeleton and $Fe_{BF}$ for the synthesis of nitrogenase, respectively.

The decomposition of nitrogenase seems to occur at night (21, 22), and therefore it is not considered during the light period in our model. Notably, nitrogenase is inhibited upon exposure to $O_2$, flowing into the pool of inactivated nitrogenase (5) at the rate [$T_{NF}^{NA}$, μmol Fe (mol C)$^{-1}$ s$^{-1}$]:

$$T_{NF}^{NA} = T_{NF_{max}}^{NA} \cdot \frac{Fe_{NF}}{Fe_{NF} + k_{Fe}^{NF}} \cdot \frac{O_2}{O_2 + k_{O_2}^{NF}}, \tag{S31}$$

where $T_{NF_{max}}^{NA}$ [μmol Fe (mol C)$^{-1}$ s$^{-1}$] is the maximal inactivation rate of nitrogenase.

The Fe in nitrogenase and photosystems from (21) were estimated from observed protein content, based on Fe atoms in per protein. PSII, Cyt *b6f*, PSI and Ferredoxin together represents photosystems. Cyt *b6f* and Ferredoxin were not measured in (21) but estimated by assuming Cyt *b6f*:PSII = 1:1 in Fe quota and Ferredoxin:PSI = 1:1 in protein content. Further details are in the supplementary information in (20).

### 8. Integration of state variables during the daytime

The diurnal change rates (basically normalized to carbon biomass) of $CH_2O$, CS, N, intracellular $O_2$ and Fe are represented in ordinary differential equations (ODEs). Note that $O_2$ is in a unit volumetric concentration (mol $O_2$ m$^{-3}$):

$$\frac{dCH_2O}{dt} = V_{CF} - V_{CS} - V_{PR} - V_{RP}, \tag{S32}$$

$$\frac{dCS}{dt} = V_{CS}, \tag{S33}$$

$$\frac{dN}{dt} = V_{NF}, \tag{S34}$$

$$\frac{dO_2}{dt} = (V_{O_2} - V_{O_2}^{PR} - V_{O_2}^{RP}) \cdot Q_C + T_{O_2}, \tag{S35}$$

$$\frac{dFe_{PS}}{dt} = T_{PS}^{BF} - T_{BF}^{PS}, \tag{S36}$$

$$\frac{dFe_{NF}}{dt} = T_{NF}^{BF} - T_{NF}^{NA}, \tag{S37}$$

$$\frac{dFe_{NF}^{NA}}{dt} = T_{NF}^{NA}, \tag{S38}$$

$$\frac{dFe_{BF}}{dt} = T_{BF}^{PSI} - T_{PSI}^{BF} \cdot + T_{NF}^{BF}, \tag{S39}$$

where $Q_C$ = 18333 mol C $m^{-3}$ is the cellular carbon biomass quota (23). ODEs are run over a12-hour light period with ode15s integrator of MATLAB (24).

### 9. Biosynthesis and growth rate

*Trichodesmium* might store newly fixed C and N during the daytime and assimilate them into biomass, mainly during the dark period (25). Therefore, for simplification, no biomass is synthesized during the light period in the model. Instead, the model calculates the amount of biomass [*Bio*, mol C (mol C)$^{-1}$] that can be synthesized using the carbohydrates, carbon skeletons and fixed N at the end of the light period. *Bio* is the smaller of N-based ($Bio_N$) and C-based biomass ($Bio_C$), with $Bio_N$ calculated by dividing fixed N to the molar N:C (0.159) (26). $Bio_C$ is calculated from the carbohydrates and carbon skeleton considering mass and energy balance. The energy needed for biosynthesis is from the respiration of carbohydrates ($CH_2O_{BIO}^{RESP}$):

$$Bio_C \cdot q_{BIO}^{ATP} \cdot (1 + \gamma_{MT}) = CH_2O_{BIO}^{RESP} \cdot q_{RESP}^{ATP}, \tag{S40}$$

where $q_{BIO}^{ATP}$ = 2 mol ATP (mol C)$^{-1}$ is the ATP requirement rate by biosynthesis (14), and $q_{RESP}^{ATP}$ = 5 mol ATP (mol C)$^{-1}$ is the ATP production rate from respiring carbohydrates (27). Meanwhile, the non-respired carbohydrates and all the carbon skeletons are involved in biosynthesis:

$$Bio_C = CH_2O - CH_2O_{BIO}^{RESP} + CS. \tag{S41}$$

$Bio_C$ then can be solved from the above two equations. Note that the carbohydrate respiration calculated in this step is counted in the daily integrated respiration as the ordinary respiration.

Noting that all the rates have been normalized to carbon biomass, *Bio* is therefore the relative increase in biomass over one day. The growth rate (*G*) is then the natural log of (1 + *Bio*) divided by 1 day.

### TABLE S1 Optimized parameters

| Symbol | Unit | Definition | Value: Low Fe (40 pM), Fixed permeability | Value: Low Fe (40 pM), Dynamic permeability | Value: High Fe (1250 pM), Fixed permeability | Value: High Fe (1250 pM), Dynamic permeability |
|---|---|---|---|---|---|---|
| $v_{RP}^{max}$ | mol C (mol C)$^{-1}$ s$^{-1}$ | Maximal respiratory protection rate | $9.6\times10^{-4}$ | $1.2\times10^{-3}$ | $9.8\times10^{-4}$ | $8.2\times10^{-4}$ |
| $T_{PS_{max}}^{BF}$ | μmol Fe (mol C)$^{-1}$ s$^{-1}$ | Maximal synthesis rate of photosystems | $5.7\times10^{-6}$ | $3.8\times10^{-6}$ | $2.0\times10^{-6}$ | $1.6\times10^{-5}$ |
| $T_{BF_{max}}^{PS}$ | μmol Fe (mol C)$^{-1}$ s$^{-1}$ | Maximal decomposition rate of photosystems | $6.3\times10^{-4}$ | $1.3\times10^{-3}$ | $1.7\times10^{-3}$ | $4.8\times10^{-3}$ |
| $T_{NF_{max}}^{BF}$ | μmol Fe (mol C)$^{-1}$ s$^{-1}$ | Maximal synthesis rate of nitrogenase | $2.2\times10^{-2}$ | $2.8\times10^{-2}$ | $2.7\times10^{-2}$ | $2.8\times10^{-2}$ |

### TABLE S2 Fixed model parameters

| Symbol | Unit | Definition | Value | Source or note |
|---|---|---|---|---|
| $v_{PET}^{max}$ | mol electron (mol C)$^{-1}$ s$^{-1}$ | Maximal PET rate | $1.0\times10^{-2}$ | This study [a] |
| $k_{Fe}^{PS}$ | μmol Fe (mol C)$^{-1}$ | Half-saturating coefficient of Fe in photosystems for PET rate | 25 | This study [a] |
| $k_{CH_2O}^{PR}$ | mol C (mol C)$^{-1}$ | Half-saturating coefficient of $CH_2O$ for photorespiration | 0.4 | This study [a] |
| $k_{O_2}^{PR}$ | mol $O_2$ m$^{-3}$ | Half-saturating coefficient of $O_2$ for photorespiration | 1.917 | This study [a] |
| $v_{CS}^{max}$ | mol C (mol C)$^{-1}$ s$^{-1}$ | Maximal production rate of CS | $8.6\times10^{-6}$ | This study [a] |
| $k_{CS}$ | mol C (mol C)$^{-1}$ | Half-saturating coefficient of CS for RP | 0.4 | This study [a] |
| $k_{CH_2O}^{CS}$ | mol C (mol C)$^{-1}$ | Half-saturating coefficient of $CH_2O$ for CS production | 0.4 | This study [a] |
| $k_{Fe_{PS}}^{PS_{syn}}$ | μmol Fe (mol C)$^{-1}$ | Half-saturating coefficient of $Fe_{PS}$ for the synthesis of photosystems | 1.0 | This study [a] |
| $k_{Fe_{PS}}^{PS_{dec}}$ | μmol Fe (mol C)$^{-1}$ | Half-saturating coefficient of $Fe_{PS}$ for the decomposition of photosystems | 25 | This study [a] |
| $k_{Fe_{BF}}^{NF_{syn}}$ | μmol Fe (mol C)$^{-1}$ | Half-saturating coefficient of $Fe_{BF}$ for the synthesis of nitrogenase | 5.0 | This study [a] |
| $T_{NF_{max}}^{NA}$ | μmol Fe (mol C)$^{-1}$ s$^{-1}$ | Maximal inactivation rate of nitrogenase | $3.3\times10^{-3}$ | This study [a] |
| $\varepsilon_{max}$ | dimensionless | Maximal relative diffusivity of cell membrane | $2\times10^{-4}$ | This study [a] |
| $k_{O_2}^{diff}$ | mol $O_2$ m$^{-3}$ | Half-saturating coefficient of $O_2$ for relative diffusivity of cell membrane | 0.213 | This study [a] |
| $N_{max}$ | mol N (mol C)$^{-1}$ | Maximal fixed storage | 0.159 | This study [b] |
| $CS_{max}$ | mol C (mol C)$^{-1}$ | Maximal CS storage | 1 | This study [c] |
| $\alpha_I$ | μmol$^{-1}$ m$^2$ s | Initial slope of $P$ versus $I$ curve | 0.01 | (14) |
| $\beta$ | (mol C)$^{-1}$ mol C s | Parameter of inhibition effect of respiration on PET | $2\times10^4$ | (7) |
| $k_{O_2}^{NF}$ | mol $O_2$ m$^{-3}$ | Half-saturating coefficient of $O_2$ for $N_2$ fixation | 0.01 | (7) |
| $d_{O_2}$ | m$^2$ s$^{-1}$ | $O_2$ diffusion coefficient at 34 PSU and 25 °C | $2.26\times10^{-9}$ | (28) |
| $k_{Fe}^{NF}$ | μmol Fe (mol C)$^{-1}$ | Half-saturating coefficient of Fe in nitrogenase for $N_2$ fixation | 91 | (20) |
| $\gamma_{MT}$ | dimensionless | Ratio of the energy consumed by maintenance to other process | 10% | (20) |
| $Fe_{TH}$ | μmol Fe (mol C)$^{-1}$ | Threshold of intracellular metabolic Fe quota | 24.4 | (20) |
| $f_{ST}$ | dimensionless | Fraction of luxury Fe uptake | 0.90 | (20) |
| | | ***Boundary conditions*** | | |
| $I_{max}$ | μmol m$^{-2}$ s$^{-1}$ | Maximal light intensity in fixed-Fe and dynamic-Fe cases under diurnally changing light intensity | 160 | |
| $O_2^E$ | mol $O_2$ m$^{-3}$ | Extracellular far-field $O_2$ | 0.213 | |
| | | ***Elemental or energy stoichiometries of metabolic activities*** | | |
| $q_{LPET}^{NADPH}$ | mol NADPH (mol electron)$^{-1}$ | NADPH/electron ratio of LPET | 0.5 | (4) |
| $q_{LPET}^{ATP}$ | mol ATP (mol electron)$^{-1}$ | ATP/electron ratio of LPET | 0.65 | (3) |
| $q_{LPET}^{O_2}$ | mol $O_2$ (mol electron)$^{-1}$ | $O_2$/electron ratio of LPET | 0.25 | (4) |
| $q_{AET}^{ATP}$ | mol ATP (mol electron)$^{-1}$ | ATP/electron ratio of MR-AET | 0.65 | (3) |
| $q_{NF}^{NADPH}$ | mol NADPH (mol N)$^{-1}$ | NADPH/N ratio of $N_2$ fixation | 3 | (9, 10) |

| | | | | |
|---|---|---|---|---|
| $q_{NF}^{ATP}$ | mol ATP (mol N)$^{-1}$ | ATP/N ratio of $N_2$ fixation | 9 | (9, 10) |
| $q_{PR}^{NADPH}$ | mol NADPH (mol C)$^{-1}$ | NADPH/C ratio of photorespiration | 4 | (8) |
| $q_{PR}^{ATP}$ | mol ATP (mol C)$^{-1}$ | ATP/C ratio of photorespiration | 7 | (8) |
| $q_{C,\,PR}^{O_2}$ | mol $O_2$ (mol C)$^{-1}$ | $O_2$/C ratio of photorespiration | 3 | (8) |
| $q_{CF}^{NADPH}$ | mol NADPH (mol C)$^{-1}$ | NADPH/C ratio of C fixation | 2 | (13) |
| $q_{CF}^{ATP}$ | mol ATP (mol C)$^{-1}$ | ATP/C ratio of C fixation | 3 | (13) |
| $q_{CCM}^{ATP}$ | mol ATP (mol C)$^{-1}$ | ATP/C ratio of CCM | 0.8 | (12) |
| $q_{BIO}^{ATP}$ | mol ATP (mol C)$^{-1}$ | ATP/C ratio of biosynthesis | 2 | (14) |
| $q_{RESP}^{ATP}$ | mol ATP (mol C)$^{-1}$ | ATP/C ratio of respiration | 5 | (27) |
| $q_{C}^{O_2}$ | mol $O_2$ (mol C)$^{-1}$ | $O_2$/C ratio of respiration | 1 | (27) |
| $Q_C$ | mol C m$^{-3}$ | Cellular carbon biomass concentration | 18333 | (23) |
| | | ***Morphological parameters of Trichodesmium*** | | |
| $L$ | m | Length of the total trichome | $554\times10^{-6}$ | (29) |
| $R$ | m | Radius of the cytoplasm | $4.80\times10^{-6}$ | (29) |
| $L_g$ | m | Thickness of cell membrane layer | 0.076 | (29) |

[a] Estimated based on model experiments under constant light intensity.

[b] By multiplying the initial C biomass with the molar N:C (0.159) of *Trichodesmium* (26).

[c] $CS_{max}$ is set to be the same as the initial C biomass.

### TABLE S3 Intermediate process and state variables

| Symbol | Unit | Definition |
|---|---|---|
| | | ***Intermediate process variables*** |
| $V_{PET}$ | mol electron (mol C)$^{-1}$ s$^{-1}$ | PET rate |
| $V_{LPET}$ | mol electron (mol C)$^{-1}$ s$^{-1}$ | LPET rate |
| $V_{AET}$ | mol electron (mol C)$^{-1}$ s$^{-1}$ | AET rate |
| $f_{AET}$ | dimensionless | Fraction of electrons in PET to AET |
| $V_{NAPDH}$ | mol NADPH (mol C)$^{-1}$ s$^{-1}$ | NADPH production rate |
| $V_{ATP}$ | mol ATP (mol C)$^{-1}$ s$^{-1}$ | ATP production rate |
| $V_{O_2}$ | mol $O_2$ (mol C)$^{-1}$ s$^{-1}$ | $O_2$ production rate |
| $V_{PR}$ | mol C (mol C)$^{-1}$ s$^{-1}$ | Photorespiration rate |
| $V_{NADPH}^{PR}$ | mol NADPH (mol C)$^{-1}$ s$^{-1}$ | NADPH consumption rate of photorespiration |
| $V_{ATP}^{PR}$ | mol ATP (mol C)$^{-1}$ s$^{-1}$ | ATP consumption rate of photorespiration |
| $V_{O_2}^{PR}$ | mol $O_2$ (mol C)$^{-1}$ s$^{-1}$ | $O_2$ consumption rate of photorespiration |
| $V_{NF}^{max}$ | mol N (mol C)$^{-1}$ s$^{-1}$ | Maximal $N_2$ fixation rate |
| $V_{NF}$ | mol N (mol C)$^{-1}$ s$^{-1}$ | $N_2$ fixation rate |
| $V_{NADPH}^{NF}$ | mol NADPH (mol C)$^{-1}$ s$^{-1}$ | NADPH consumption rate of $N_2$ fixation |
| $V_{ATP}^{NF}$ | mol ATP (mol C)$^{-1}$ s$^{-1}$ | ATP consumption rate of $N_2$ fixation |
| $V_{ATP}^{CCM}$ | mol ATP (mol C)$^{-1}$ s$^{-1}$ | ATP consumption rate of CCM |
| $V_{NADPH}^{CF}$ | mol NADPH (mol C)$^{-1}$ s$^{-1}$ | NADPH consumption rate of C fixation |
| $V_{ATP}^{CF}$ | mol ATP (mol C)$^{-1}$ s$^{-1}$ | ATP consumption rate of C fixation |
| $V_{CS}$ | mol C (mol C)$^{-1}$ s$^{-1}$ | Carbon skeleton production rate |
| $V_{RP}$ | mol C (mol C)$^{-1}$ s$^{-1}$ | Respiratory protection rate |
| $V_{O_2}^{RP}$ | mol $O_2$ (mol C)$^{-1}$ s$^{-1}$ | $O_2$ consumption rates by respiratory protection |
| $\varepsilon$ | dimensionless | Relative diffusivity of cell membrane |
| $T_{O_2}$ | mol $O_2$ m$^{-3}$ s$^{-1}$ | $O_2$ diffusion rate between cytoplasm and ambient environment |
| $Fe_M$ | µmol Fe (mol C)$^{-1}$ | Intracellular Fe quota in metabolism |
| $Fe_{ST}$ | µmol Fe (mol C)$^{-1}$ | Intracellular Fe quota in storage |
| $T_{PS}^{BF}$ | µmol Fe (mol C)$^{-1}$ s$^{-1}$ | Synthesis rate of photosystems |
| $T_{BF}^{PS}$ | µmol Fe (mol C)$^{-1}$ s$^{-1}$ | Decomposition rate of photosystems |
| $T_{NF}^{BF}$ | µmol Fe (mol C)$^{-1}$ s$^{-1}$ | Synthesis rate of nitrogenase |
| $T_{NF}^{NA}$ | µmol Fe (mol C)$^{-1}$ s$^{-1}$ | Inactivation rate of nitrogenase |
| $Bio$ | mol C (mol C)$^{-1}$ | New synthesized biomass |
| $Bio_N$ | mol C (mol C)$^{-1}$ | New synthesized N-based biomass |
| $Bio_C$ | mol C (mol C)$^{-1}$ | New synthesized C-based biomass |
| $CH_2O_{BIO}^{RESP}$ | mol C (mol C)$^{-1}$ | Respired carbohydrates to fulfill the energy need for biosynthesis |
| $G$ | d$^{-1}$ | Specific growth rate |
| | | ***State variables*** |
| $CH_2O$ | mol C (mol C)$^{-1}$ | Carbohydrate |
| $CS$ | mol C (mol C)$^{-1}$ | Carbon skeleton |
| $N$ | mol N (mol C)$^{-1}$ | Fixed N |
| $O_2$ | mol $O_2$ m$^{-3}$ | Intracellular $O_2$ |
| $Fe_{PS}$ | µmol Fe (mol C)$^{-1}$ | Fe in photosystems |
| $Fe_{NF}$ | µmol Fe (mol C)$^{-1}$ | Fe in active nitrogenase |
| $Fe_{NF}^{NA}$ | µmol Fe (mol C)$^{-1}$ | Fe in inactivated nitrogenase |
| $Fe_{BF}$ | µmol Fe (mol C)$^{-1}$ | Fe in buffer |

Note: The initial values (t = 0) of $CH_2O$, $CS$ and $N$ are set to be 0, and initial $O_2$ concentration is the same as that of ambient $O_2$ (0.213 mol $O_2$ m$^{-3}$).

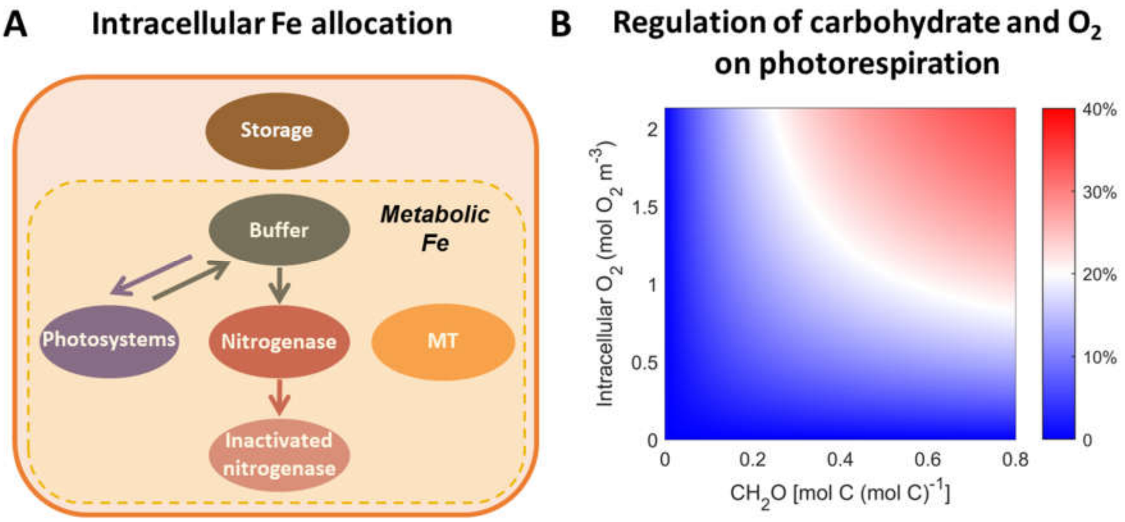


**FIG S1 (A) Model structure of intracellular Fe allocation in *Trichodesmium* and (B) regulation of carbohydrate and $O_2$ on photorespiration.** (**A**) The model encompasses fundamental intracellular Fe pools, including storage and five metabolic pools (photosystems, active and inactivated nitrogenase, maintenance, buffer and storage). Note that all inactivated nitrogenase by $O_2$ does not decompose and Fe in it is not recycled during model period (daytime). (**B**) The regulation effect is represented by the ratio of $V_{PR}$ to $V_{PR}^{max}$.

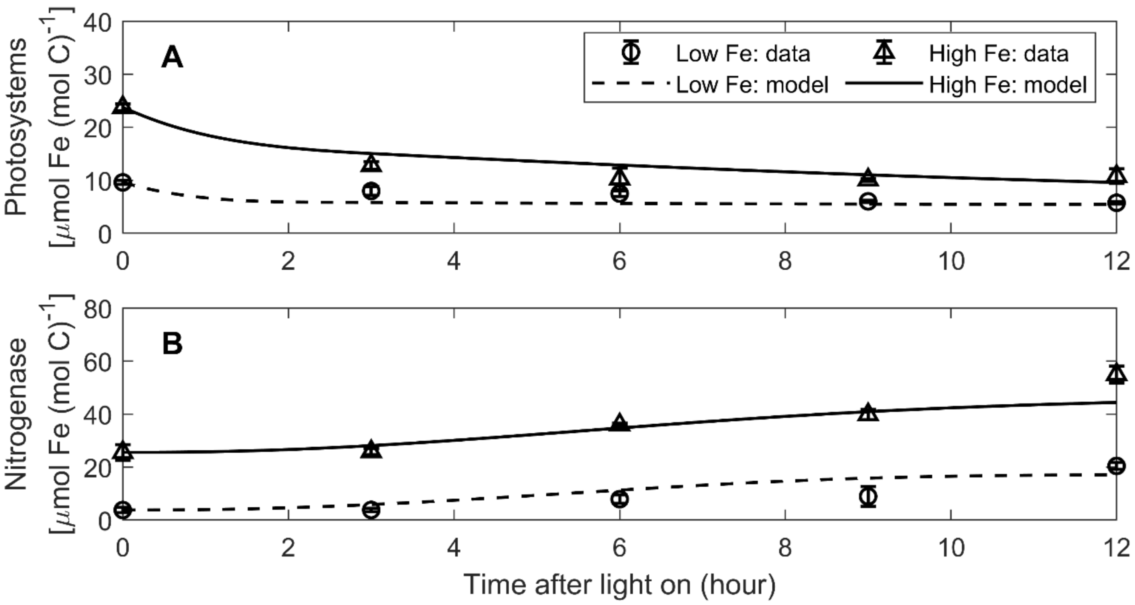


**FIG S2 Modeled and observed diurnal variations of Fe in photosystems and nitrogenase.** The observational data are from (21). The model is simulated with dynamic $O_2$ permeability of the cell membrane under both low-Fe and high-Fe conditions. Light intensity in the model is diurnally constant as that in (21). Error bars represent one standard deviation.

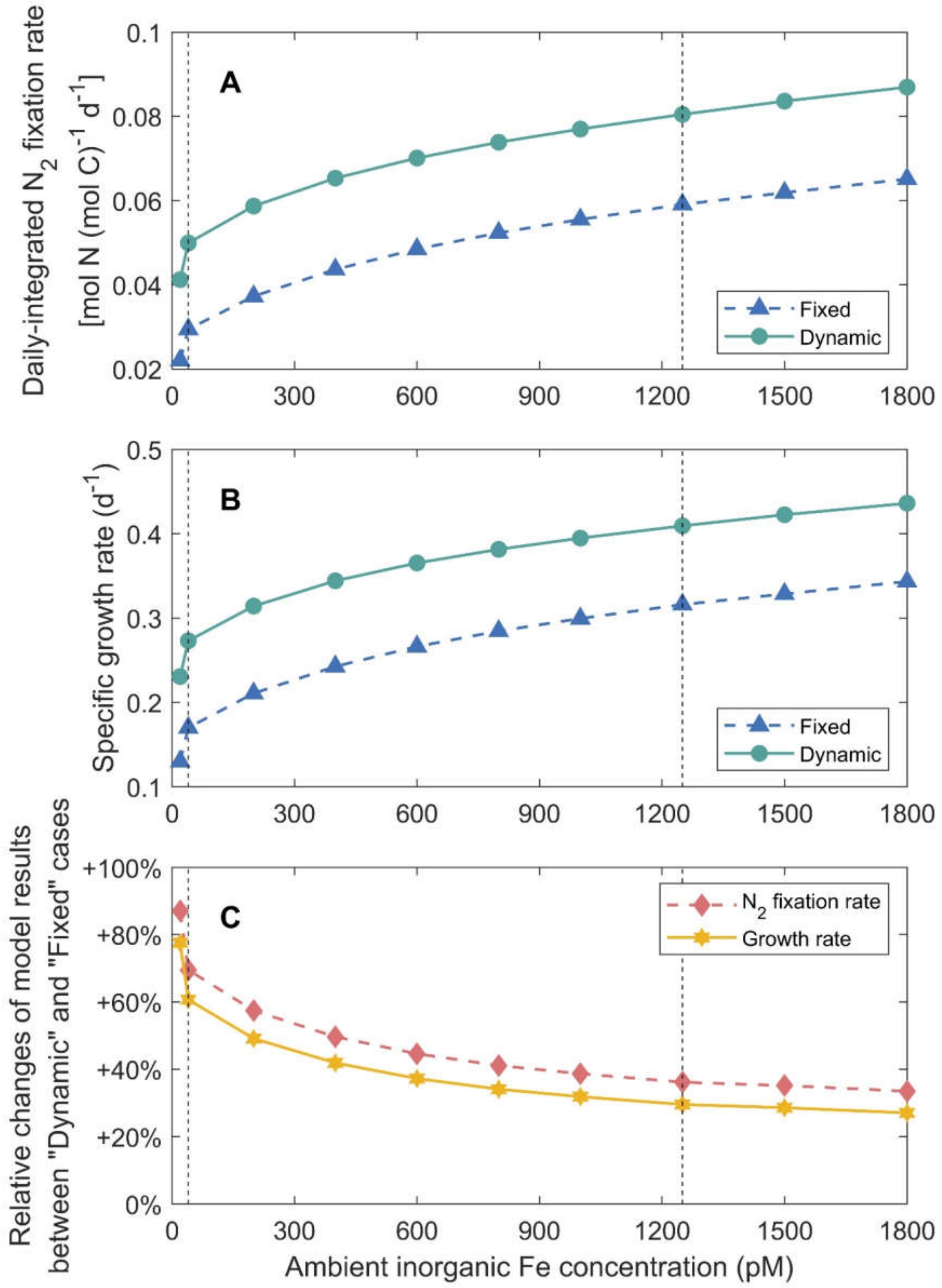


**FIG S3 Simulated daily-integrated $N_2$ fixation rates (A), growth rates (B) and their relative changes (C) between "Dynamic permeability" and "Fixed permeability" cases under different ambient inorganic iron concentrations.** Black dashed lines represent ambient inorganic iron concentrations (40 and 1250 pM) used in the standard model cases.

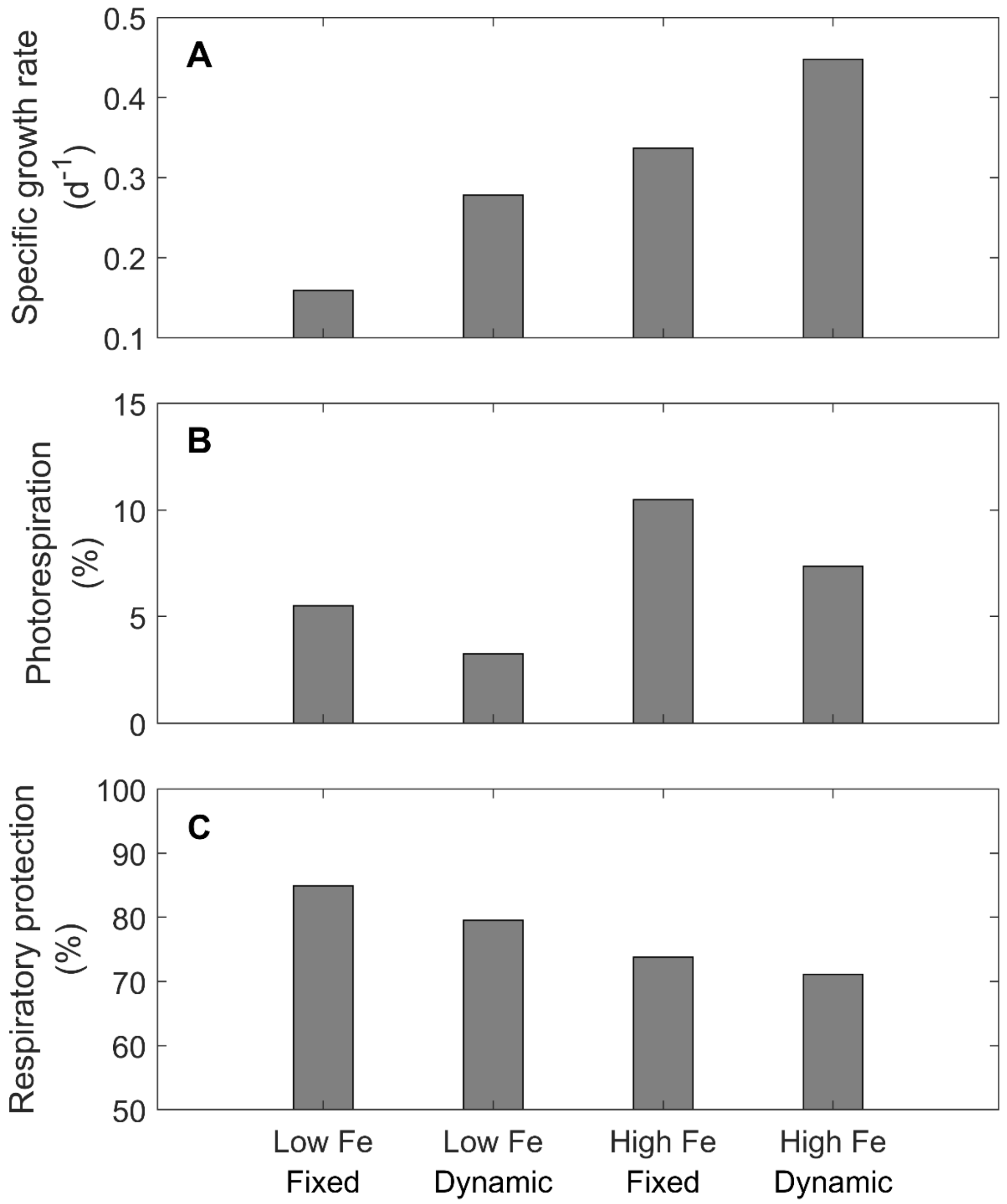


**FIG S4 Results of model experiments under the constant light intensity, including growth rates (A), gross fixed carbon allocated to respiratory protection (B) and photorespiration (C).** The model is simulated with diurnally fixed or dynamic $O_2$ permeability of the cell membrane under low-Fe (40 pM) and high-Fe (1250 pM) conditions.

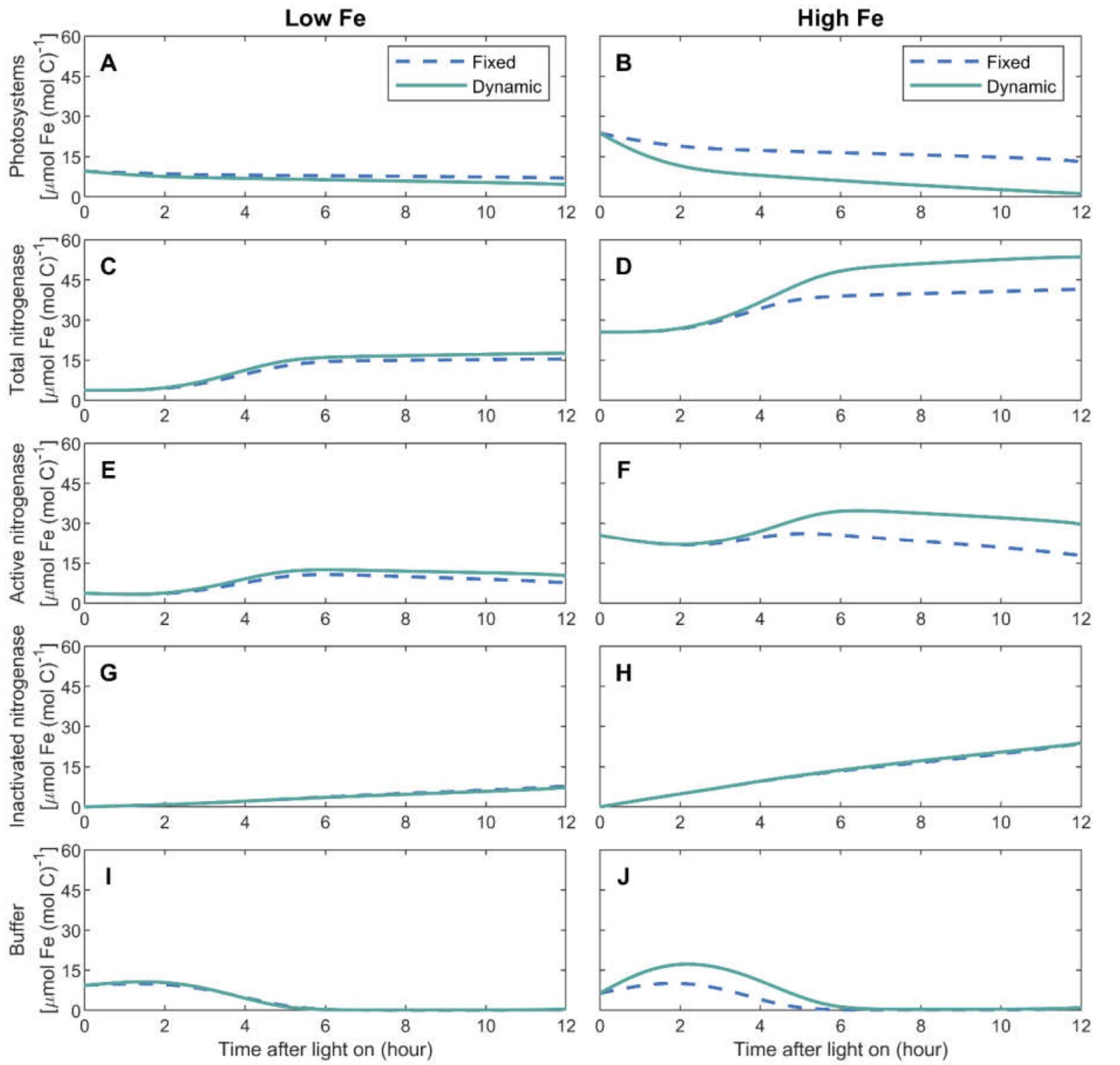


**FIG S5 Simulated diurnal variations of Fe in photosystems, total, active and inactivated nitrogenase, and buffer.** The model is simulated with diurnally fixed or dynamic $O_2$ permeability of the cell membrane under low-Fe (40 pM) (**A**, **C**, **E**, **G** and **I**) and high-Fe (1250 pM) (**B**, **D**, **F**, **H** and **J**) conditions.
